# Goblet cell invasion promotes breaching of respiratory epithelia by an opportunistic human pathogen

**DOI:** 10.1101/2023.08.13.553119

**Authors:** A. Leoni Swart, Benoît-Joseph Laventie, Rosmarie Sütterlin, Tina Junne, Xiao Yu, Evdoxia Karagkiozi, Rusudan Okujava, Urs Jenal

## Abstract

While commensal bacteria generally respect natural barriers of the human body, pathogens are able to breach epithelia, invade deeper tissue layers and cause life-threatening infections. *Pseudomonas aeruginosa*, an opportunistic human pathogen, is a leading cause of severe hospital-acquired pneumonia, with mortality rates as high as 50% in mechanically ventilated patients^1–3^. Effective colonization and breaching of lung mucosa are hallmarks of *P. aeruginosa* pathogenesis^4^. Although virulence factors and behavioral strategies of *P. aeruginosa* have been described^5,6^, it has remained unclear how this pathogen disseminates on functional mucosal surfaces, how it avoids mucociliary clearance and how it invades the tissue barrier. Using fully differentiated human lung epithelia, we demonstrate that *P. aeruginosa* efficiently spreads on the apical tissue surface before it breaches epithelia by specifically invading mucus secreting goblet cells. Internalization leads to host cell death and expulsion and the formation of ruptures of the epithelial barrier. Rupture sites are rapidly colonized by extracellular bacteria through active chemotaxis, leading to increasing tissue damage and successful pathogen translocation to the unprotected basolateral side of the epithelium. We show that cell invasion is promoted by two Type-6 toxin secretion systems (T6SS), while Type-3 (T3SS) mediates cell death of infected goblet cells. T3SS mutants invade goblet cells normally, but internalized bacteria fail to trigger goblet cell expulsion and instead show unrestrained intracellular replication. While the effective shedding of infected host cells reveals potent tissue protection mechanisms, the discovery of an intracellular lifestyle of *P. aeruginosa* in human lung epithelia provides new entry points into investigating the intersection of antibiotic and immune mechanisms during lung infections. By demonstrating that *P. aeruginosa* uses a combination of specific virulence factors and collective behavior to invade goblet cells and breach the lung tissue barrier from within, these studies reveal novel mechanisms underlying lung infection dynamics under physiological conditions.

## Introduction

Organs of the human respiratory, digestive and urogenital tracts are lined by a single layer of epithelial cells, which serves as an effective physical and biological barrier against invading pathogens. Mucosal epithelia are monolayers of highly polarized cells that are interconnected through tight junctions, multiprotein complexes that regulate barrier permeability. Secretion of a mucus layer represents a first line of defense by limiting the contact between luminal pathogens and the tissue surface. Epithelia also harbor receptors sensing the presence of bacteria on the apical surface and staging a powerful immune response, while receptors for pathogen adherence and internalization are generally sequestered to the basolateral side of the membrane^7^. Accordingly, pathogens preferentially adhere to areas of epithelial damage to cross the barrier and reach the basal membrane compartment^8,9^.

The human respiratory tract is a complex organ system facilitating the exchange of oxygen and carbon dioxide. Its large environmental-epithelial interface is continuously exposed to toxins, irritants, and pathogens. The pseudostratified epithelium of the upper airways contains multi-ciliated cells, mucus-secreting goblet cells, neuroendocrine cells and basal cells, which secrete surfactant and serve as resident stem cells for self-renewal during homeostasis and after injury (**Fig. 1a**). Inhaled microbes are actively removed through mucociliary clearance or by pulmonary innate immune defenses^1,2,10,11^. Nevertheless, the lung remains the main entry portal for airborne pathogens, with respiratory infections representing a major cause of death worldwide and the highest health burden of years lost through death or disability^12,13^. For example, with 1.4 million deaths each year tuberculosis is the most common lethal infectious disease next to COVID-19^14^. Moreover, critical priority pathogens as defined by the WHO, like *Pseudomonas aeruginosa*, *Acinetobacter baumannii* and *Klebsiella pneumoniae*, are among the dominant agents of acute and chronic respiratory infections, causing pneumonia or sepsis and leading to severe inflammatory damage, acute lung injury and inflammation or acute respiratory distress syndrome^2,3^.

**Fig. 1.**
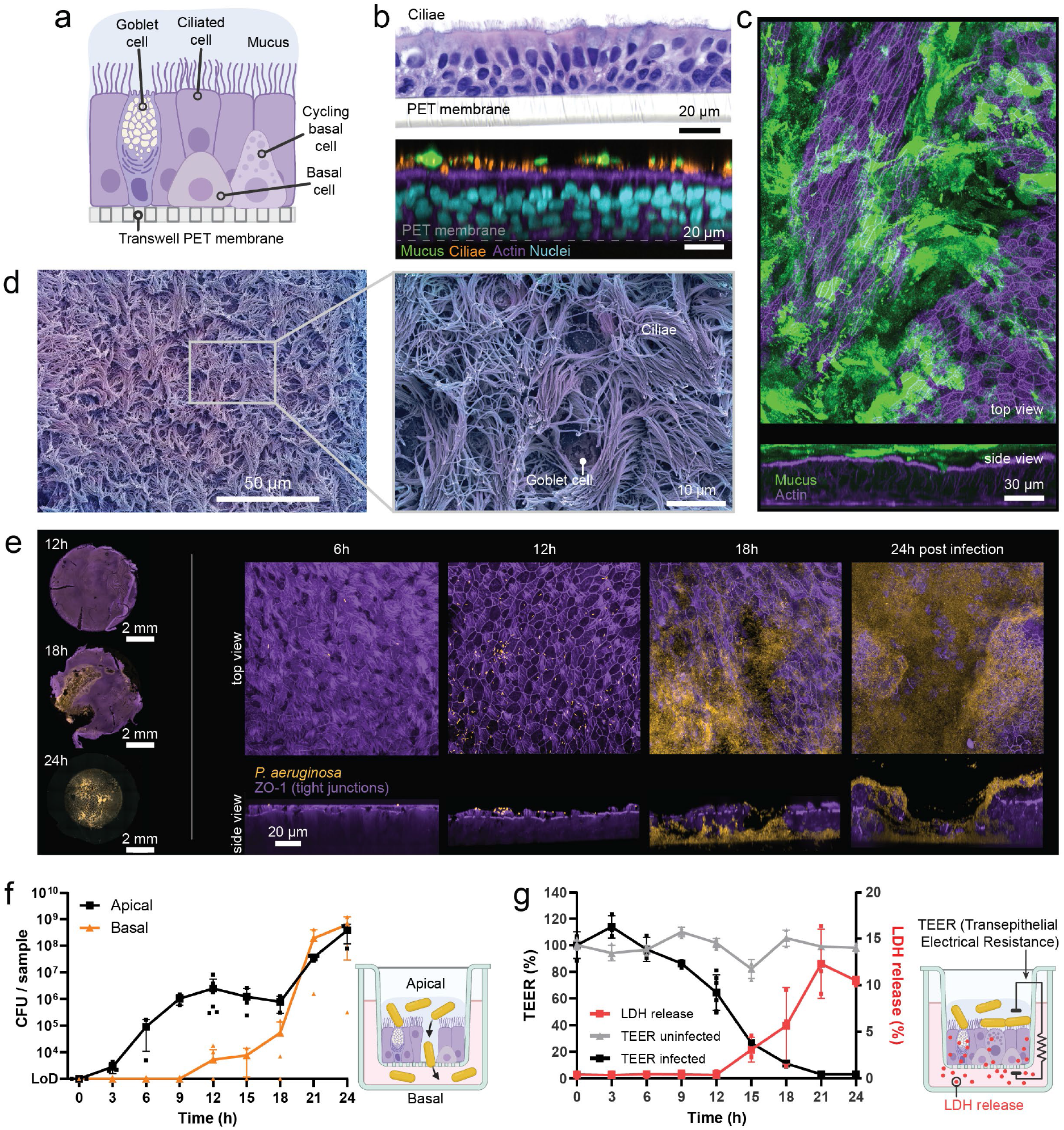
Human airway model exposes individual steps of *P. aeruginosa* lung infection. **a.** Schematic of the ALI upper airway model architecture and cellular composition. **b.** Histology (H&E staining) and Immunocytochemistry images of fully differentiated airway epithelium after 30 days of air exposure. **c.** Mucus secreted on the apical surface of the airway epithelium. **d.** Upper airway epithelium surface imaged using SEM. **e**. ALI lung model colonization by *P. aeruginosa* (ICC). **f.** *P. aeruginosa* population on the airway epithelium (black) and in the basal compartment after tissue breaching (orange). **g.** Epithelial barrier integrity (TEER) and tissue viability (LDH release) during the infection. For all panels: Inoculum=10^3^, *n*=3. ICC staining: bacteria: constitutive chromosomal expression of mNeonGreen, nuclei: DAPI, mucus: Muc5AC, membrane: CellMask, cilia: acetylated-α-tubulin, actin: phalloidin-647, tight junctions: ZO-1 Alexa Fluor 555.

Here, we use *P. aeruginosa*, an opportunistic human pathogen, to dissect the key steps of human lung tissue colonization. *P. aeruginosa* is a leading cause of severe hospital-acquired pneumonia, with mortality rates as high as 50% for mechanically ventilated patients^1^. *P. aeruginosa* is also the leading cause of respiratory morbidity and mortality in patients with cystic fibrosis^10^ and a frequent cause of exacerbations in individuals with advanced chronic obstructive pulmonary disease^15^. While many of the virulence factors and behavioral strategies of *P. aeruginosa* have been identified, their exact role in infecting human lung mucosa is not well-understood^5,16^. Understanding how *P. aeruginosa* colonizes and breaches human lung mucosa may unravel new pathogen vulnerabilities and therapeutic options, which are increasingly limited due to the continued spread of multi-drug resistant strains.

Studies with monolayers of non-differentiated lung cell lines indicated that the apical surface of the lung epithelium is refractory to bacterial toxin injection, arguing that epithelial cell polarity contributes to the defense against *P. aeruginosa* infection^17,18^. Accordingly, *P*. *aeruginosa* was reported to use a paracellular transmigration route, exploiting altered cell-cell junctions at sites of cell division, tissue damage or when senescent cells are expelled from the cell layer^19^. Others have proposed a transcellular migration route, in which internalized bacteria transform a sub-domain of the apical cell surface into a basolateral-like domain^20,21^. Because these studies were generally performed with cell cultures or monolayers of epithelial cell lines lacking fully differentiated tissue barrier function, the underlying infection mechanisms have remained controversial^22,23^. Likewise, animal models often lack the experimental accessibility to investigate infection processes with high temporal and spatial resolution and their predictive value for disease phenotypes has remained controversial^24–26^.

The recent development of organoids, self-organizing 3D structures grown from stem cells that recapitulate essential aspects of organ structure and function, provides robust models to study infections directly in fully differentiated human tissues^27^. Originally established for gastrointestinal tissue^28^, several models are now available, including bladder^29^, pancreas^30^, liver^31^ as well as upper^32^ and lower^33^ airways. Lung organoids have been utilized to model specific disease phenotypes^34,35^ and to study infections by respiratory viruses or parasites^36,37^. Here, we use human upper airway tissues to visualize and quantify *P. aeruginosa* lung infection with unprecedented spatial and temporal resolution. Our studies provide a detailed mechanistic frame for how human pathogens overcome the mucus barrier and rapidly spread on mucosal tissue and how they combine invasion of specialized cell types and collective behavior to rapidly and effectively breach the barrier function of the lung epithelium.

## Results

### Human airway model exposes individual steps of *P. aeruginosa* lung infection

To study *P. aeruginosa* infections of intact human lung epithelia, we employed an upper airway tissue model. Bronchial epithelial progenitor cells were expanded and subsequently cultivated on PET membrane inside Transwell cell culture inserts. Confluent cell layers were then exposed to air for a minimum of 24 days, stimulating differentiation into a pseudostratified mucociliary epithelium that closely mimics human lung tissue (**Fig. 1a,b**; Extended data **Fig. 1a-e**). Flow cytometry analysis and immunocytochemistry (ICC) confirmed the presence of all major cell types, including ciliated, goblet and basal cells (**Fig. 1a**, Extended data **Fig. 2a,b**)^38^. Pseudo-stratified tissues were fully functional, secreting a confluent mucus layer and exposing highly active cilia on their surface, coordinately beating with frequencies typically observed in human bronchia (**Fig. 1c,d**, Extended data **Fig. 2c**, Extended data **Movie 1**)^11^. Epithelial barrier function was confirmed by trans-epithelial electrical resistance (TEER) measurements (Extended data **Fig. 2d**) and the presence of tight junctions throughout the tissue (Extended data **Fig. 2e**). Tissue functionality, viability and regeneration was maintained for up to 60 days (Extended data **Fig. 1a-e**, Extended data **Fig. 2c,d,f**, Extended data **Movie 2**).

**Fig. 2.**
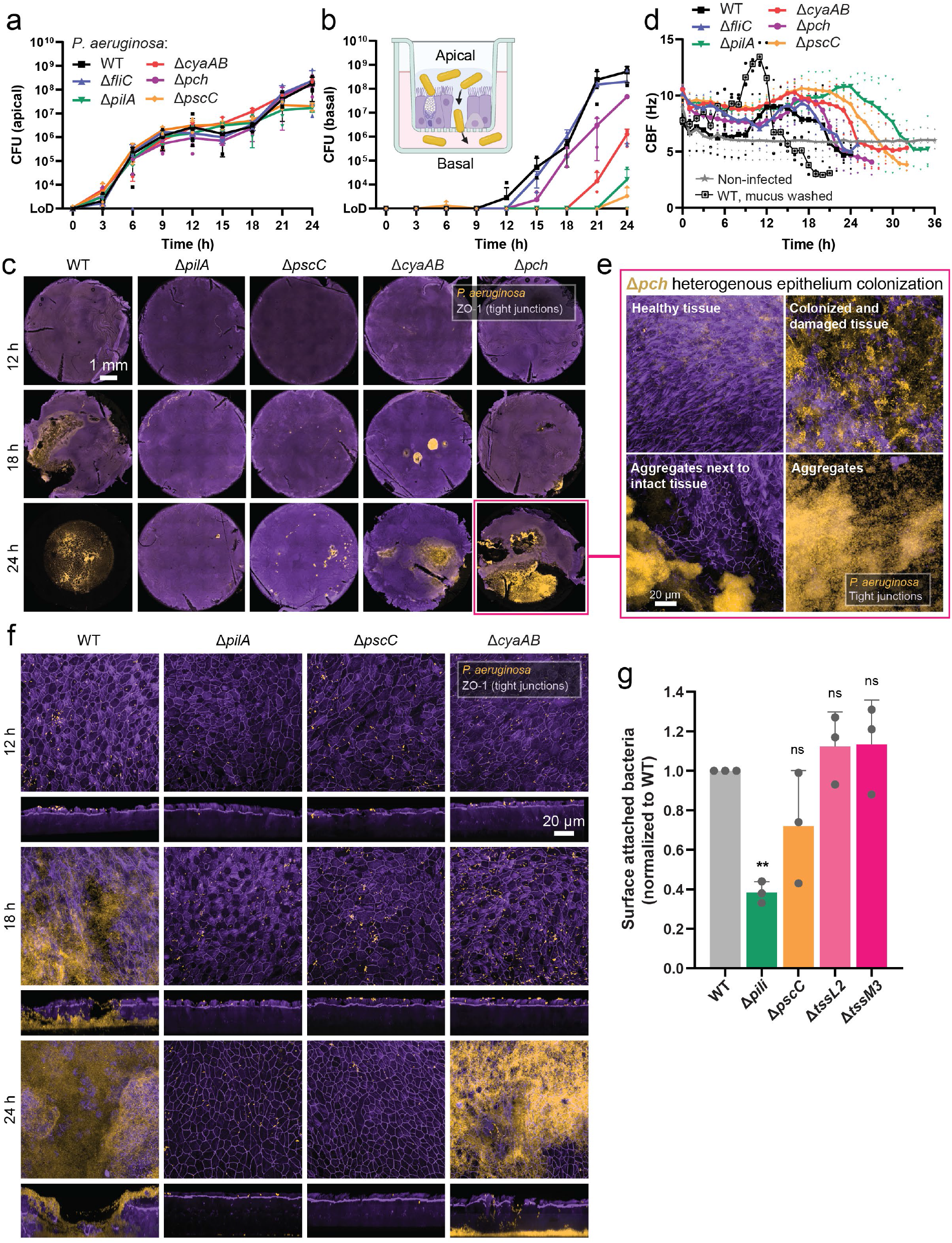
Specific virulence factors promote *P. aeruginosa* lung tissue colonization and breaching. **a, b**. *P. aeruginosa* wild type and mutant strains population on the airway epithelium (**a**) and in the basal compartment after tissue breaching (**b**). **c, f.** Colonization of airway epithelia by *P. aeruginosa* wild type and mutants. 10x overview of the entire transwell (**c**) and 100x details (**f**) (ICC). **d**. Cilia beating frequency of the airway epithelium during *P. aeruginosa* infection measured by live microscopy. **e.** The impaired spreading of the Δ*pch* strain leads to a considerable spatial heterogeneity of colonization and tissue damage (ICC, 24 h post infection). **g**. Quantification of the airway epithelium colonization by *P. aeruginosa* mutants from ICC images. Dunnett’s One-Way ANOVA compared to WT; ** p<0.01; ns: not significant. For all panels: Inoculum=10^3^, *n*=3. ICC staining: bacteria: constitutive chromosomal expression of mNeonGreen, tight junctions: ZO-1 Alexa Fluor 555.

Inoculation of lung tissues with *P. aeruginosa* on the air side revealed a characteristic and highly reproducible bi-phasic behavior, with two phases of rapid growth separated by an extended lag-phase of about 10 hours (**Fig. 1e,f**). During the initial phase of infection, bacteria rapidly replicated in the mucus layer, with few bacteria being in close proximity to the epithelial surface (**Fig. 1e**). Mucus is constantly produced by goblet cells and removed by the mucociliary escalator, thereby forming an effective physical barrier that prevents pathogenic microbes from reaching the surface of the epithelium^39^. In line with its protective function, removal of the mucus layer prior to infection accelerated tissue breaching and destruction (Extended data **Fig. 3a,b**). At 12 hours post-infection, the tissue surface was densely colonized with non-replicating *P. aeruginosa* bacteria that started to infect epithelial cells, as indicated by a steady increase of apoptosis, a loss of barrier integrity and tissue viability, and by bacteria gradually breaking through to the basal compartment (**Fig. 1e-g**; Extended data **Fig. 3c**; Extended data **Movie 3**). At 18 hours post-infection, tissue integrity was strongly compromised at multiple sites with bacteria being detected predominantly at the basolateral side of the epithelial layer. This coincided with the onset of the second phase of exponential growth and loss of tissue integrity and viability (**Fig. 1e-g**; Extended data **Fig. 3d**). 24 hours post-infection, the tissue was completely destroyed with the remaining fragments of lung tissue being covered by *P. aeruginosa* (**Fig. 1e**; Extended data **Fig. 3e**). Replacing the cell culture medium in the basolateral compartment with phosphate buffer did not alter infection kinetics, excluding the possibility that bacterial growth was mediated by nutrients provided by the cell culture medium (Extended data **Fig. 3f**). Not unexpectedly, infection kinetics was dependent on the inoculum size, with bi-phasic replication being progressively concealed and tissue destruction accelerated at increasing initial pathogen loads (Extended data **Fig. 3g-i**). These experiments provided the frame to investigate *P. aeruginosa* infection of human lung tissue in greater detail.

**Fig. 3.**
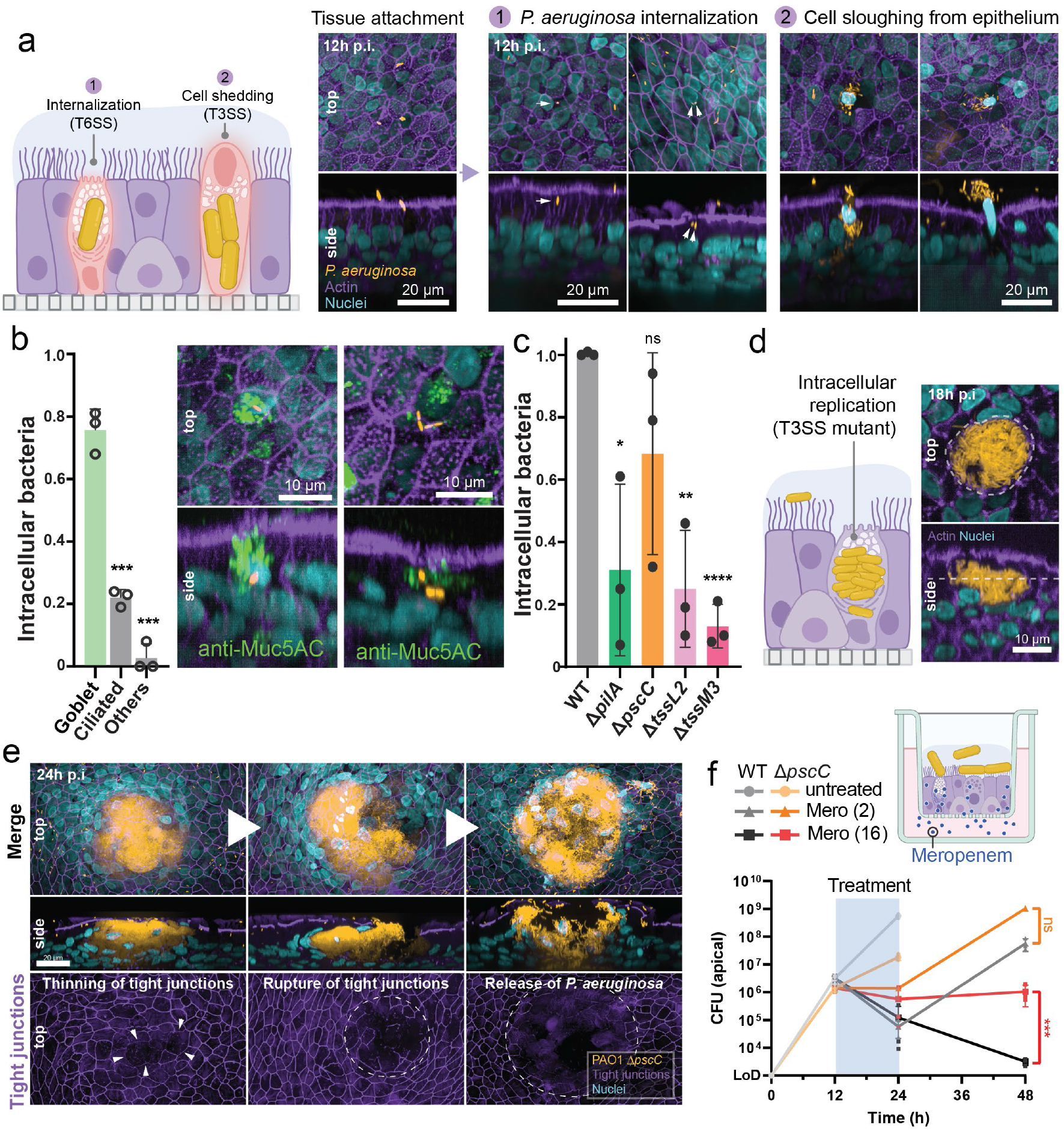
Goblet cells serve as entry gates for *P. aeruginosa* invasion of lung epithelia. **a.** Schematic and microscopy images of *P. aeruginosa* tissue attachment, **(1)** internalization into epithelial cells and **(2)** subsequent cell sloughing. **b**. Quantification of specific cell types with internalized *P. aeruginosa* and corresponding microscopy example. Unpaired t test compared to Goblet; *** p<0.0002. **c**. Quantification of internalized *P. aeruginosa* wild type and mutants in epithelial cells. Unpaired t test compared to WT; * p<0.0332; ** p<0.0021; *** p<0.0002; ns: not significant. **d**, **e**. Schematics (**d**) and microscopy images (**e**) of intracellular replication by the T3SS-mutant (Δ*pscC*). Continued replication of the T3SS-mutant eventually leads to thinning and rupture of the tight junction and release of *P. aeruginosa*. **f**. Meropenem treatment of airway epithelia infected with *P. aeruginosa* wild type or T3SS mutant (Δ*pscC*). Antibiotics (2 or 16 µg/ml) were added and removed at t12 and t24 p.i., respectively. Unpaired t test; *** p<0.0002; ns: not significant. For all panels: Inoculum=10^3^, *n*=3. ICC staining: bacteria: constitutive chromosomal expression of mNeonGreen, nuclei: DAPI, mucus: Muc5AC, actin: phalloidin-647, tight junctions: ZO-1 Alexa Fluor 555.

### Specific virulence factors promote *P. aeruginosa* lung tissue colonization and breaching

*P. aeruginosa* harbors a range of virulence factors, including endo- and exotoxins, adhesins like Type IV pili (T4P), as well as type-III (T3SS) and type-VI (T6SS) toxin secretion systems^40^, most of which are tightly regulated and coordinated during colonization of host surfaces^41–45^. Moreover, to disseminate on host surfaces *P. aeruginosa* utilizes motors like rotary flagella or dynamic T4P^46,47^. Motility and virulence are tightly regulated and coordinated during surface colonization by a cascade of second messengers, including cAMP and c-di-GMP^48–52^. While cAMP promotes virulence gene expression on surfaces^43,49^, c-di-GMP directs a bimodal behavioral program that promotes *P. aeruginosa* virulence through asymmetric divisions producing a motile and an adhesive offspring^48^. Because *P. aeruginosa* virulence behavior is generally studied on artificial surfaces or with cell culture systems lacking tight barrier function, we set out to test the role of known virulence factors in lung tissue colonization and breaching. Mutants lacking the flagellum (Δ*fliC*), T4P (Δ*pilA*), or T3SS (Δ*pscC*), as well as mutants unable to synthesize cAMP (Δ*cyaAB*) or to maintain c-di-GMP asymmetry during cell division (Δ*pch*) all showed normal growth in an axenic medium mimicking the human lung environment^53^ and during the initial stage of lung tissue infection (**Fig. 2a**; Extended data **Fig. 4a**). However, Δ*cyaAB*, Δ*pilA* or Δ*pscC* mutants showed greatly delayed tissue breaching, emphasizing their key importance in tissue invasion and cytotoxicity (**Fig. 2b,c**; Extended data **Fig. 4b,c**). This effect was prominently observed when monitoring cilia beating frequencies, which increased temporarily in response to *P. aeruginosa* infection followed by a rapid drop during pathogen invasion and tissue destruction (**Fig. 2d**)^54^. Tissues infected with Δ*cyaAB*, Δ*pilA,* Δ*pch* or Δ*pscC* mutants did not show increased cilia beating, but maintained active cilia for extended periods (**Fig. 2d**; Extended data **Fig. 4d**).

**Fig. 4.**
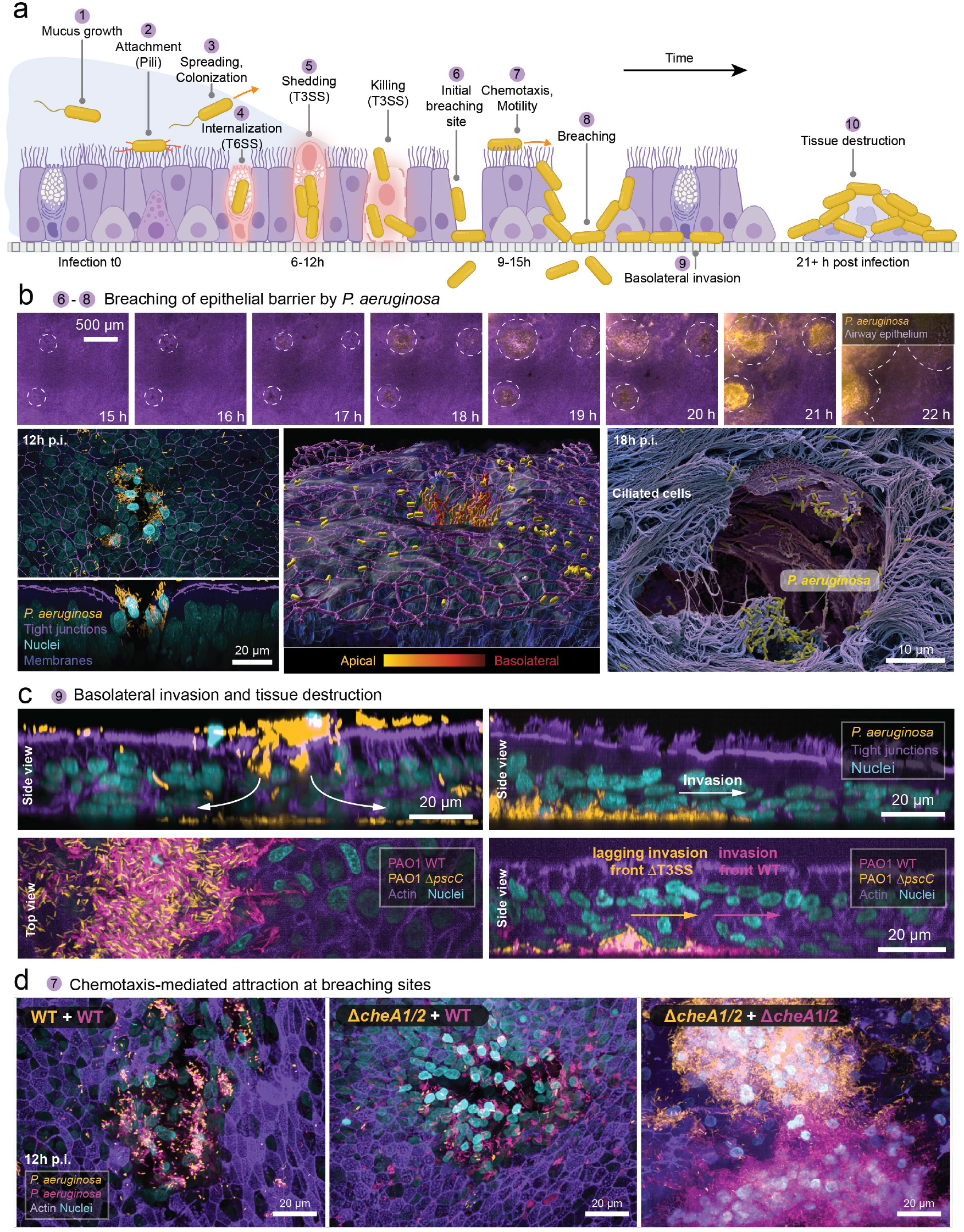
Pathogen-mediated local tissue destruction provides access to the basolateral tissue side. **a.** Schematic of the different phases of *in vitro* lung tissue colonization by *P. aeruginosa* PAO1 wild type. **b**. Stills of low magnification (10x, upper panels) live imaging visualizing initiation and expansion of tissue breaching events (Movie S2). Detailed images (100x ICC, lower left and middle panels or SEM, lower right panel) of *P. aeruginosa* breaching sites. **c**. Invasion of the basolateral side of the epithelium by *P. aeruginosa* after establishing a breaching site. Initiation of basolateral invasion requires a functional T3SS (lower panels). **d**. Breaching site colonization in samples infected 1:1 with distinctly labeled (pink and yellow) wild type strains (left panel) or Δ*cheA*1/2 mutant strains (right panel) or wild type and Δ*cheA*1/2 mutant strains (middle panel). For all panels: Inoculum=10^3^, *n*=3. Staining: bacteria: constitutive chromosomal expression of mNeonGreen or mCherry, nuclei: DAPI, mucus: Muc5AC, membrane: CellMask, actin: phalloidin-647, tight junctions: ZO-1 Alexa Fluor 555.

Intriguingly, a Δ*pch* mutant, although displaying only a mild infection delay (**Fig. 2a,b**; Extended data **Fig. 4c,d**), showed strongly compromised dissemination with large parts of the tissue remaining sterile and entirely intact for up to 24 hrs (**Fig. 2c,e**). Thus, the ability of *P. aeruginosa* to divide asymmetrically to generate motile and adherent offspring strongly promotes tissue colonization. Tissue spreading could in principle engage flagellar or T4P-based motors. *P. aeruginosa* lacking a flagellum showed a delayed tissue response and failed to adopt the typical arrangement at early infection stages with single bacteria evenly spread-out on the tissue surface. Instead, the mutant clustered in small aggregates, indicative of its inability to effectively spread on the lung tissue surface (Extended data **Fig. 4e**). Defining a clear role for T4P in tissue spreading is more difficult, as these organelles serve both as primary adhesins and as surface motors^48^. Accordingly, *P. aeruginosa* lacking T4P (Δ*pilA*) showed strongly impaired attachment to lung epithelia (**Fig. 2f,g**). Thus, the asymmetric program generates motile and sessile individuals that cooperate to promote *P. aeruginosa* dissemination and virulence on epithelial surfaces.

### Goblet cells serve as entry gates for *P. aeruginosa* invasion of lung epithelia

Upon depleting the mucus layer, *P. aeruginosa* started interfering with the epithelial cell layer (**Fig. 3a**; Extended data **Fig. 5a**). Tissue imaging revealed frequent pathogen internalization by host cells in the upper layer of the epithelium at 12 hours post-infection, a time point coinciding with the first indications of host tissue response, *e.g.*, increased cilia beating, but preceding cytotoxicity and tissue damage. While many host cells contained single internalized bacteria, others harbored multiple pathogens, indicating that *P. aeruginosa* is able to proliferate inside differentiated human lung epithelial cells (**Fig. 3a**; Extended data **Fig. 5b**). This was supported by infections with a mixture of *P. aeruginosa* strains expressing different fluorophores, which exclusively revealed internalized bacteria of the same color (Extended data **Fig. S5c**). Strikingly, almost 80% of host cells with internalized pathogens were goblet cells (**Fig. 3b**; Extended data **Fig. 5d**), even though they only represent a fraction of the epithelial cell population (Extended data **Fig. 2b**)^55^. Internalization events were frequent, with roughly 1% of all surface exposed host cells being invaded by *P. aeruginosa* during the initial breaching phase.

It is possible that *P. aeruginosa* actively targets and invades mucus secreting goblets cells once nutrients in the mucus layer are depleted. This view is supported by the observation that mutants lacking T4P showed reduced internalization (**Fig. 3c**). Moreover, mutants lacking H2-T6SS (Δ*tssL2*) and H3-T6SS (Δ*tssM3*), two toxin delivery systems known to promote *P. aeruginosa* internalization into non-phagocytic cells^56,57^, showed strongly reduced numbers of infected goblet cells. In contrast, goblet cell invasion was only marginally reduced upon infection with mutants lacking the T3SS (Δ*pscC*) (**Fig. 3c**). However, while goblet cells infected with *P. aeruginosa* wild type were gradually sloughed into the apical lumen by the lung tissue and rapidly consumed by extracellular pathogens (**Fig. 3a**; Extended data **Fig. 6a,b**), internalized T3SS mutants showed no cytotoxicity and continued to divide inside goblet cells leading to large intracellular bacterial aggregates (**Fig. 3d**). This led to a massive volume increase of infected host cells, physically straining the tissue to an extent that eventually resulted in thinning and rupturing of tight junctions, cell bursting, pathogen release and spreading to neighboring cells (**Fig. 3e**; Extended data **Fig. 7a,b**). This explains why T3SS mutants are still able to breach lung epithelia, although with a strong time delay as compared to *P. aeruginosa* wild type (Extended data **Fig. 4b,d**). Moreover, the extended intracellular lifestyle of T3SS mutants effectively protects against antibiotic therapy, as treatment of infected lung tissues with high concentrations of meropenem eradicated *P. aeruginosa* wild type but failed to clear the T3SS mutant (**Fig. 3f**).

Based on these experiments and based on the key role of T3SS in host cell cytotoxicity and tissue breaching (**Fig. 2b**), we propose that *P. aeruginosa* actively invades goblet cells and that internalized pathogens cause massive host cell death and sloughing in a process that critically depends on the secretion of toxic effectors through the T3SS. While the molecular details of goblet cell invasion remain to be disclosed, we propose that mucus secreting cells serve as specific entry gates, allowing *P. aeruginosa* to bypass the natural barrier functions of the lung epithelium.

### Pathogen-mediated local tissue destruction provides access to the basolateral tissue side

The above experiments indicated that *P. aeruginosa* breaches the lung epithelial barrier using a combination of T6SS-mediated goblet cell invasion and T3SS-mediated cytotoxicity (**Fig. 4a**). Because we did not observe uniform tissue breakdown following initial cell invasion, we assume that most events were resolved by cell detachment and tissue repair processes^58^. However, starting 14 hours post-infection, macroscopic cavities emerged in the lung tissue at much lower frequency, gradually enlarging over time and rapidly accumulating increasing numbers of *P. aeruginosa* bacteria (**Fig. 4b**; Extended data **Fig. 8a-c**; Extended data **Movie 2**). Based on their frequency, their highly synchronous appearance and close temporal succession to goblet cell invasion, we postulate that tissue cavities are caused by a combination of massive T3SS-induced cytotoxicity of mucus-producing cells and gradually overwhelmed tissue repair functions. While tight junctions had remained largely intact during the initial invasion of goblet cells, they rapidly disintegrated at tissue cavity sites, indicating that at this stage of infection the lung tissue suffers from local destruction, while the surrounding lung tissue remained intact (**Fig. 4b**). Thus, the observed cavities represent specific passageways through which *P. aeruginosa* overcomes the epithelial barrier and rapidly migrates to the basal side of the tissue. Accordingly, pathogens accumulated on the basolateral surface of the tissue in close proximity of breaching sites, followed by their rapid lateral spread, leading to complete tissue destruction over the next few hours (**Fig. 4c**, Extended data **Fig. 8b**). Infections with a 1:1 mixture of *P. aeruginosa* wild type and T3SS mutant labeled with different fluorophores revealed that the T3SS may also be critically important for this final step of the infection process. Although the T3SS mutant was able to co-colonize breaching sites generated by *P. aeruginosa* wild type, mutant bacteria were never observed at the leading edge of the basolateral invasion wave expanding away from the original breaching sites (**Fig. 4c**). Thus, gradual pathogen expansion at the basolateral side of the epithelia likely requires T3SS-mediated cytotoxicity.

### Collective pathogen behavior promotes tissue breaching

We noticed that during the formation of tissue cavities, *P. aeruginosa* cell numbers rapidly increased at sites of tissue breaching, leading to a gradual expansion of the population deeper into the tissue (**Fig. 4c**, Extended data **Fig. 8b**). While this could result from a gradual replication of a founder clone driven by the release of fresh nutrients from lysed host cells, efficient host cell desquamation likely limits the overall efficiency of such a strategy. Alternatively, a directed and highly cooperative behavior of the pathogen, with bacteria rapidly moving in from distant locations on the tissue surface to sites of tissue damage, could greatly accelerate this process. To distinguish between these two possibilities, we infected lung tissues with a mixture of *P. aeruginosa* expressing different fluorophores, respectively. We expected that clonal expansion would result in homogenously colored populations at breaching sites, while an active assembly of invading communities would generate mixed color populations. The observation that all breaching sites contained balanced mixtures of differentially colored bacteria strongly argued for the latter (**Fig. 4d**, Extended data **Fig. 9**). Similarly, mixed infections with wild type and a mutant unable to perform chemotaxis (Δ*cheA1*/*2*) produced mixed populations at breaching sites, indicating that chemotacting bacteria can invade breaching sites generated by peers that are unable to sense chemical gradients. In contrast, differentially labeled chemotaxis mutants generated homogenous clonal populations at each breaching site (**Fig. 4d**, Extended data **Fig. 9**). These findings demonstrate that *P. aeruginosa* uses its chemosensory apparatus to reach sites of tissue damage, arguing that the invading pathogen population engages on a highly cooperative behavior to rapidly overcome tissue defenses and secure access to the basolateral surface of the tissue barrier.

## Discussion

Using fully differentiated and functional epithelia of the upper respiratory tract we have uncovered that the opportunistic human pathogen *P. aeruginosa* makes use of a sophisticated mechanisms to colonize and breach human lung tissue barriers. In a first phase of infection, *P. aeruginosa* rapidly expands by exploiting nutrients available in the mucus layer^59^. Only once the mucus is depleted and pathogen growth has stalled, *P. aeruginosa* attacks the underlaying epithelial tissue. We find that tissue penetration is initiated by the specific invasion of goblet cells, the major secretory cells in the superficial epithelium of the large airways. Although it is possible that goblet cell selectivity is the result of pathogen sampling by these secretory cells^60^, our observations suggest that this process is driven by the pathogen itself. Two of the three T6S systems of *P. aeruginosa*, H2-T6SS and H3-T6SS, are involved in goblet cell invasion, both of which are known to promote *P. aeruginosa* uptake into epithelial cells by translocating effector proteins that interfere with the PI3K/Akt pathway and with the tubulin ring complex^21,56,57,61^. Hence, T6SS-mediated rearrangement of the cytoskeletal network promotes entry of *P. aeruginosa* into goblet cells, a process that marks the initial step of epithelial invasion and crossing.

Because intact tight junctions effectively protect lung epithelia^62^, *P. aeruginosa* was shown to preferentially infect damaged tissue sites to access the less protected basolateral side of the barrier^8,9,63^. Consequently, interactions of *P. aeruginosa* with the apical surface of intact lung epithelia are rare and limited to transient cell extrusion events^64^. However, large-scale invasion of goblet cells offers an effective mechanism to bypass the tissue barrier, providing the pathogen with direct access to internal nutrients and to less protected tissue surfaces. If goblet cells play a role as ‘trojan horses’ in *P. aeruginosa* infections, the T3SS has an important and specific function in ‘opening the city gate’. We found that T3SS is not required for goblet cell entry but mediates intracellular cytotoxicity and goblet cell expulsion. Thus, the pathogen uses T3SS as a ‘bottle opener’ to stage its attacks from within the lung tissue or at sites of goblet cell extrusion, where basolateral surfaces are being transiently exposed before cell polarity and tight junctions are fully established during tissue repair processes^58^. This is in line with recent findings that *P. aeruginosa* can persist in vacuoles of corneal and bronchial human epithelial cells but escapes to the cytoplasm in a T3SS-dependent manner^65^. Mutants lacking T3SS showed massively delayed cytotoxicity and failed to develop the larger tissue cavities, linking tissue breaching and pathogen propagation directly to T3SS-mediated goblet cell cytotoxicity. Although the details of T3SS-mediated cytotoxicity within goblet cells remain to be investigated, a prominent intracellular role for T3SS could help explain why monoclonal antibodies targeting T3SS have failed to reduce *P. aeruginosa* nosocomial pneumonia incidence in *P. aeruginosa* colonized mechanically ventilated patients^66,67^. Moreover, intracellular growth of *P. aeruginosa* during lung tissue infection may explain the observed selective loss of T3SS and other virulence factors during chronic infections^68,69^. *P. aeruginosa* lacking T3SS activity can freely replicate intracellularly, accumulating large internalized pathogen subpopulations. It is possible that intracellular *P. aeruginosa* pathogens are resilient to clearance by the immune system and/or antibiotic therapy, forming a protected reservoir that drives chronic infections in Cystic Fibrosis patients^70^. SARS-CoV2 also preferentially infects lung goblet cells and increased SARS-CoV-2 replication in bronchial epithelia of COPD patients was linked to COPD-associated goblet cell hyperplasia^71^. This raises the possibility that goblet cell hypertrophy or hyperplasia a typically feature of COPD or CF patients^72–75^ also contribute to increased susceptibility to *P. aeruginosa* infections.

Finally, our experiments revealed that *P. aeruginosa* overcomes the lung tissue barrier as a population of cooperating individuals. Through an asymmetric division *P. aeruginosa* engages on classical ‘division of labor’ by simultaneously generating motile and surface adherent cells to optimize infection and dissemination. Similar cooperative mechanisms were shown to stabilize virulence programs^76,77^, diversify metabolic capacities^78–80^ or modulate stress tolerance during infections^81–83^. Furthermore, the motile population cooperates by performing chemotaxis, thereby accelerating colonization of original tissue rupture sites presumably by responding to nutrients released from dead host cells. Chemotaxis is known to promote pathogen colonization of host tissue sites^84–89^ and may well be of critical importance for *P. aeruginosa* to accelerate the infection process in the presence of host immune cells. To obtain a more realistic view on *P. aeruginosa* infection kinetics of human lung epithelia will require adding immune cells to the tissue models and to investigate the balance between pathogen colonization and host immune response. This will help uncover novel pathogen components driving epithelial colonization and breaching, inform on the specific host tissue responses, and provide important entry points to investigate the intersection of antibiotic and immune mechanisms.

## Acknowledgements

Authors are thankful to Julie Sollier for critical reading and editing of the manuscript; Sara Roig, Kai Schleicher, Alexia Loynton-Ferrand and Laurent Guerard (Biozentrum - Imaging core facility), Janine Bögli (Biozentrum - FACS core facility), Carola Alampi and Mohamed Chami (Biozentrum – BioEM facility) for technical support; Marek Basler for strains; and Irina Schuler for help with histology preparations. This study was supported by the Swiss National Science Foundation NCCR AntiResist (51NF40_180541 to U.J.) and financial support from the Roche Institute of Translational Bioengineering (ITB).

## Author contributions

Conceptualization: A.L.S, B.-J.L., R.O., U.J. Methodology: A.L.S, B.-J.L., R.S., T.J., X.Y., E.K., R.O., R.O., U.J. Formal analysis: A.L.S, B.-J.L., R.O., U.J. Investigation: A.L.S, B.-J.L., R.S., T.J., X.Y., E.K., R.O. Writing: original draft from A.L.S, B.-J.L., U.J. Funding acquisition: R.O., U.J. Supervision: R.O., U.J.

## Data availability

The data that support the findings of this study will be openly available in the public repository Zenodo (a link and DOI will be provided ahead of publication). The microscopy datasets generated and analyzed during the current study are available from the corresponding author on reasonable request.

## Extended data

**Extended data Fig. 1.**
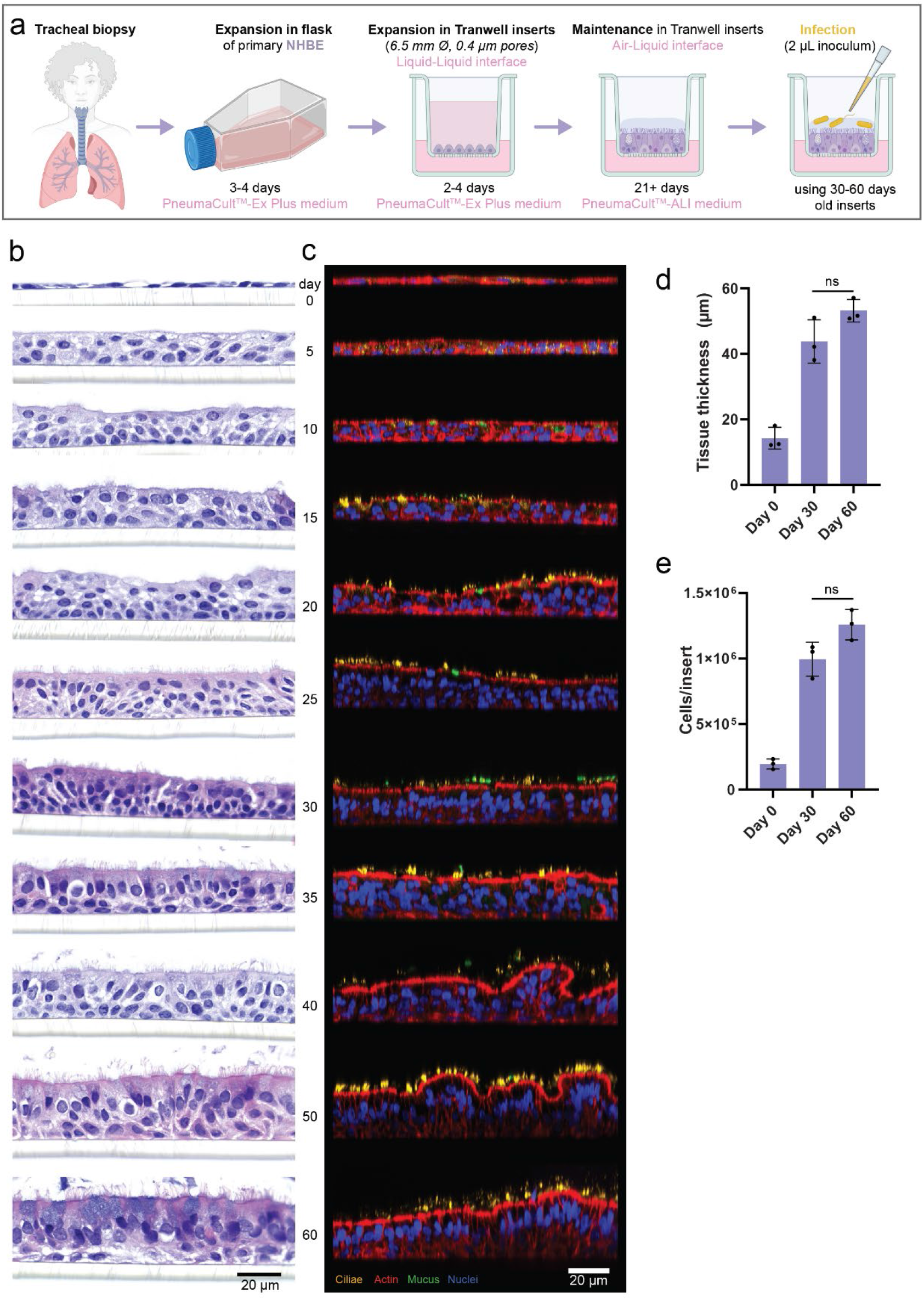
Development of *in vitro* lung tissues. **a.** Schematics of the cell culture process to develop fully differentiated (+21 days post airlift) airway epithelium on Transwell inserts and subsequent apical infection. **b**, **c**. Development of the lung tissue post airlift (day 0 to 60) visualized by histology with hematoxylin and eosin staining (**b**) or ICC (**c**). ICC staining: nuclei: DAPI, mucus: Muc5AC, actin: phalloidin-647, cilia: acetylated-α-tubulin. **d**, **e**. Quantification of the thickness (**d**) and total cell numbers (**e**) of tissues at day 0, 30 or 60 post airlift, inferred from ICC images. For (**d**) and (**e**): Statistical significance was calculated using Kruskal–Wallis *H*-test with Dunnett’s multiple comparison test. ns; not significant. For all panels: *n*=3.

**Extended data Fig. 2.**
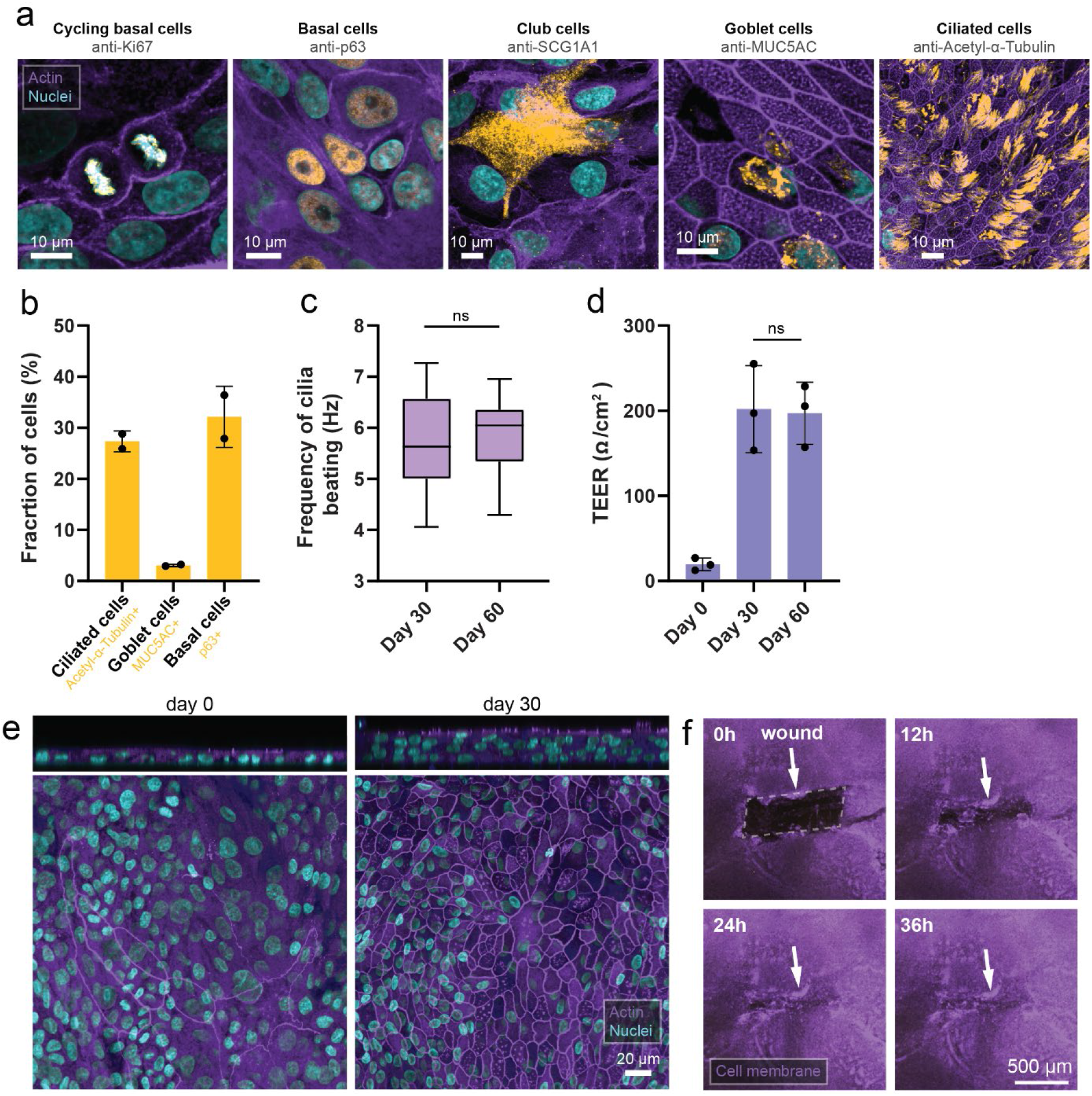
Cellular composition and functional properties of the bronchial lung epithelium model. **a**, **b**. Visualization (**a**) and quantification (**b**) of the major cells types present in the bronchial epithelium by ICC or FACS, respectively. *n*=2. **c.** Measurements of cilia beating frequencies by microscopy at day 30 or 60 post airlift. *n*=36 inserts; Box plot: center line, median; box limits, upper and lower quartiles; whiskers, min-max. **d.** Assessment of epithelial barrier integrity (TEER) of lung epithelium at day 0, 30 or 60 post airlift. *n*=3. **e**. Visualization of tight junctions by ICC of lung epithelium at day 0 or 30 post airlift. For (**d**) and (**e**): Statistical significance was calculated using Kruskal–Wallis *H*-test with Dunnett’s multiple comparison test. ns; not significant. **f.** Wound healing after physical tissue damage (indicated by the white arrow) of the lung epithelium was followed by time-lapse live imaging for 36 hours (see: Extended data Movie S2). Staining: nuclei: DAPI, cycling basal cells: mucus: Muc5AC, cycling basal cells: α-Ki67, basal cells: α-p63, club cells: α-SCG1A1, cilia: acetylated-α-tubulin, actin: phalloidin-647, tight junctions: ZO-1 Alexa Fluor 555, membrane: CellMask.

**Extended data Fig. 3.**
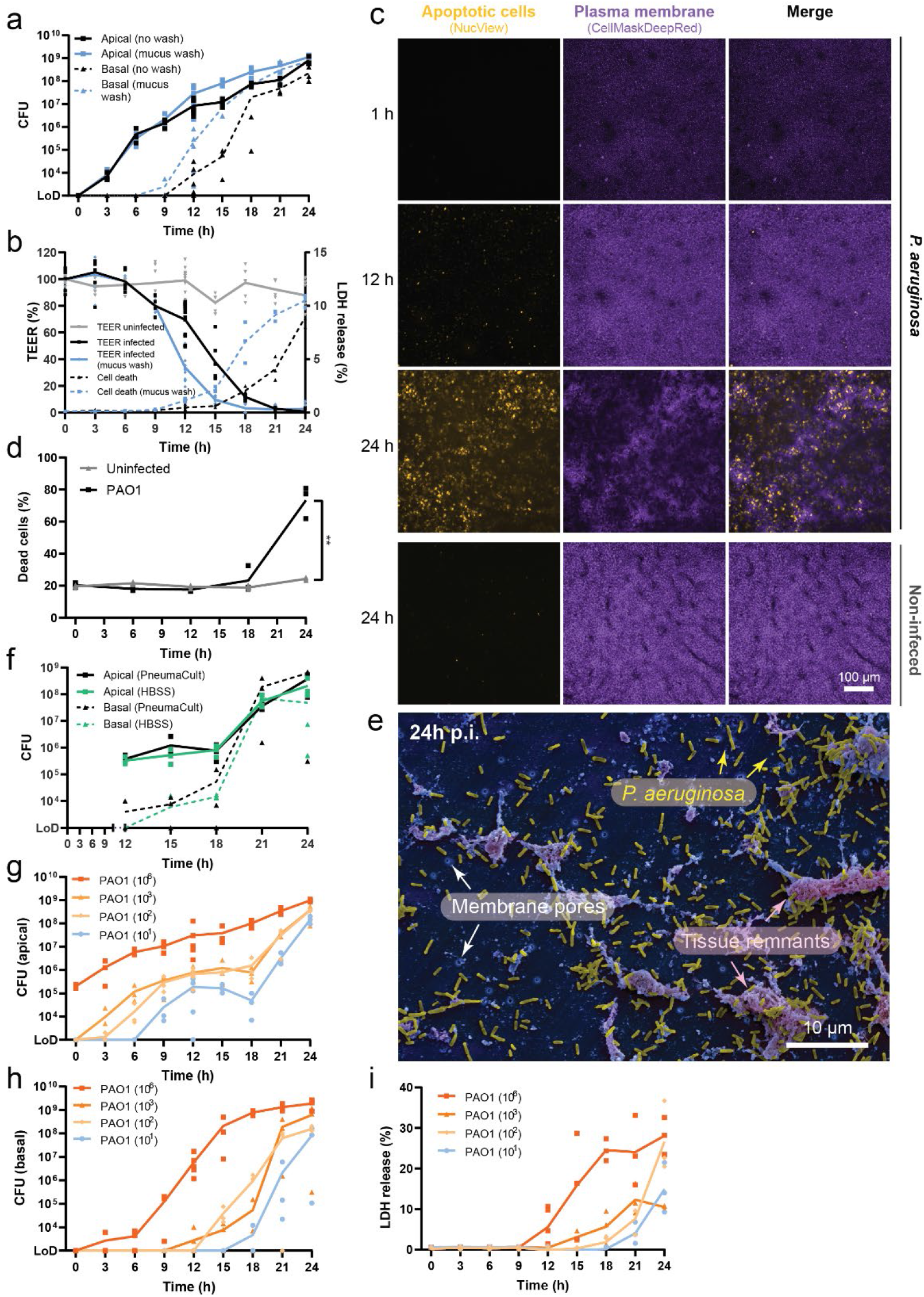
Human airway model exposes distinct steps of *P. aeruginosa* tissue colonization. **a.** *P. aeruginosa* growth on the apical side of the lung tissues (continuous line) and spread to the basal compartment after tissue breaching (dotted line) with (black) and without (blue) apical mucus upon infection. **b.** Epithelial barrier integrity (TEER) and tissue viability (LDH release) during infection in absence (blue) or presence (black) of mucus. **c**. Stills of live imaging (25x) of apoptotic events upon infection with *P. aeruginosa*. Staining: apoptotic cells: NucView (caspase-3/7 activity), membrane: CellMask **d**. Quantification of dead epithelial cells by FACS, using a viability marker (Zombie NIR), upon infection with *P. aeruginosa*. **e**. Complete tissue destruction by *P. aeruginosa* PAO1 wild type recorded by SEM. **f**. *P. aeruginosa* population on the airway epithelium (continuous line) and in the basal compartment (dashed line) with PneumaCult-ALI culture medium (black) or HBSS buffer (green) in the basolateral compartment. **g**, **h**, **i**. *P. aeruginosa* population on the apical (**g**) and basal side (**h**) of the lung tissues as well as tissue viability (LDH release) (**i)** upon infection with different inoculum sizes as indicated. For panels a-d: Inoculum=10^3^. For all panels *n*=3.

**Extended data Fig. 4.**
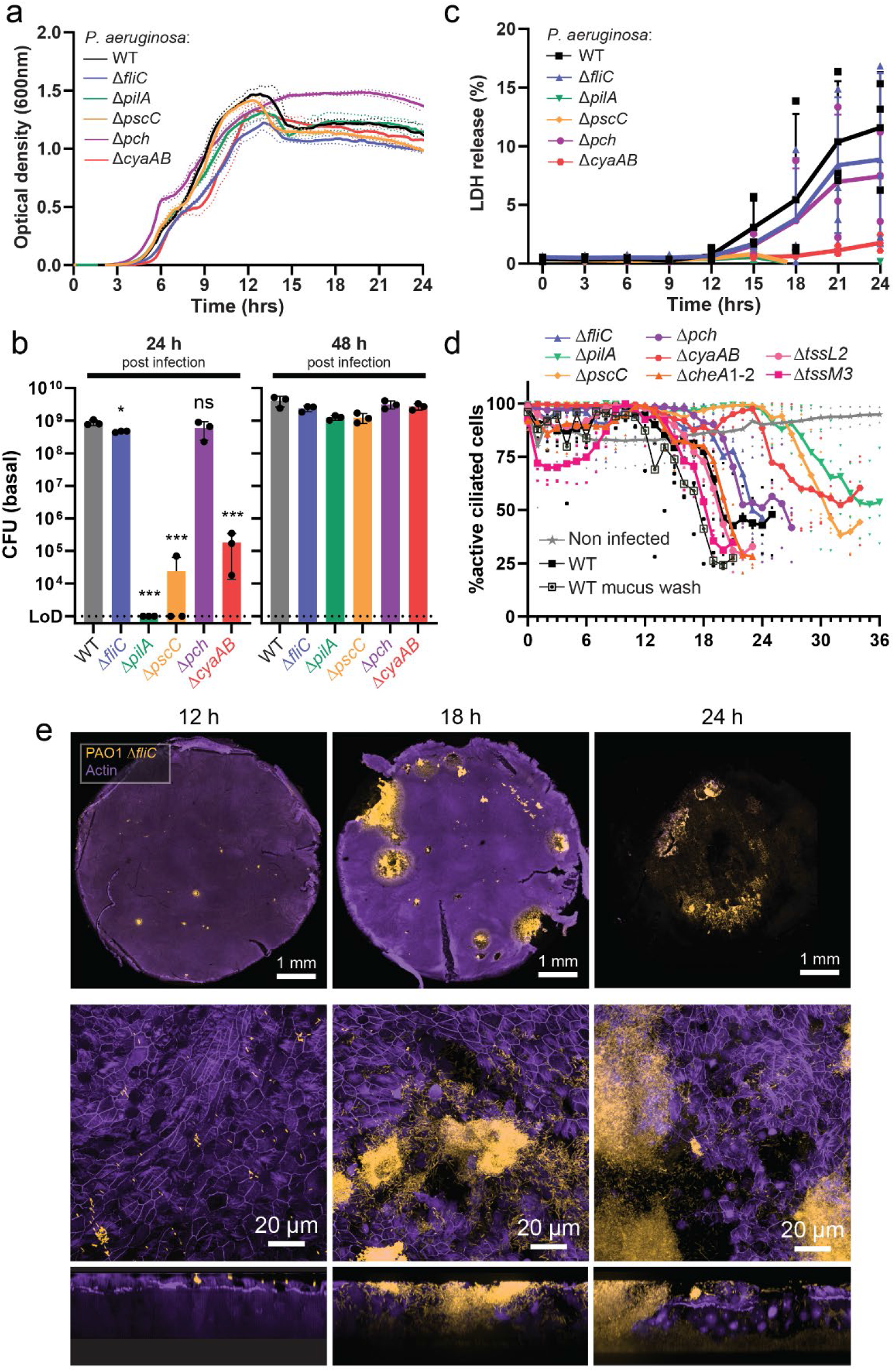
Virulence factors promoting *P. aeruginosa* lung tissue colonization and breaching. **a**. Growth of *P. aeruginosa* wild type and mutant strains in SCFM medium^1^. **b**. *P. aeruginosa* population in the basolateral compartment up to 48 h p.i. Inoculum=10^3^, *n*=3. Dunnett’s One-Way ANOVA compared to WT; * p<0.1; *** p<0.001; ns: not significant. **c**. tissue viability (LDH release) during infection with *P. aeruginosa*. **d**. Percentage actively beating cilia (>3.5Hz) of the airway epithelium during *P. aeruginosa* infection measured by live microscopy. **e**. *P. aeruginosa* Δ*fliC* colonization of the airway epithelium. 10x overview of the entire transwell (upper panel) and 100x details (middle and lower panels) (ICC). Staining: nuclei: DAPI, actin: phalloidin-647. For all panels: Inoculum=10^3^. *n*=3.

**Extended data Fig. 5.**
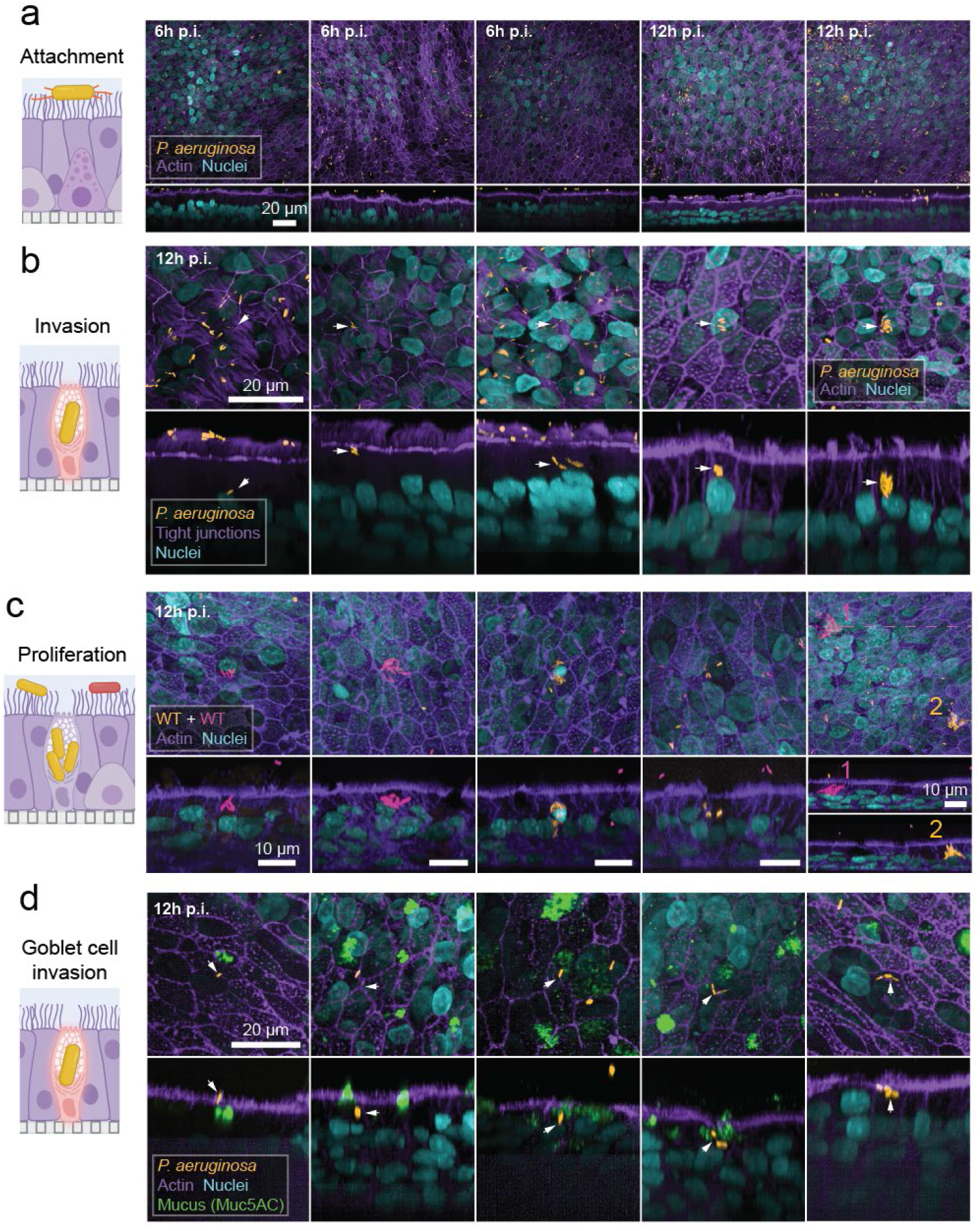
Selective internalization into goblet cells allows *P. aeruginosa* to invade lung epithelia. **a-d.** Representative examples of 100x ICC images of *P. aeruginosa* attaching to and invading lung epithelial cells at 6 h and 12 h p.i. Attached (**a**), internalized (**b**) and replicating bacteria (**c**) are shown. Specificity goblet cell invasion is illustrated in (**d**) using an additional stain for mucus. For all panels: Inoculum=10^3^. *n*=3. Staining: bacteria: constitutive chromosomal expression of mNeonGreen, nuclei: DAPI, mucus: Muc5AC, membrane: CellMask, actin: phalloidin-647, tight junctions: ZO-1 Alexa Fluor 555.

**Extended data Fig. 6.**
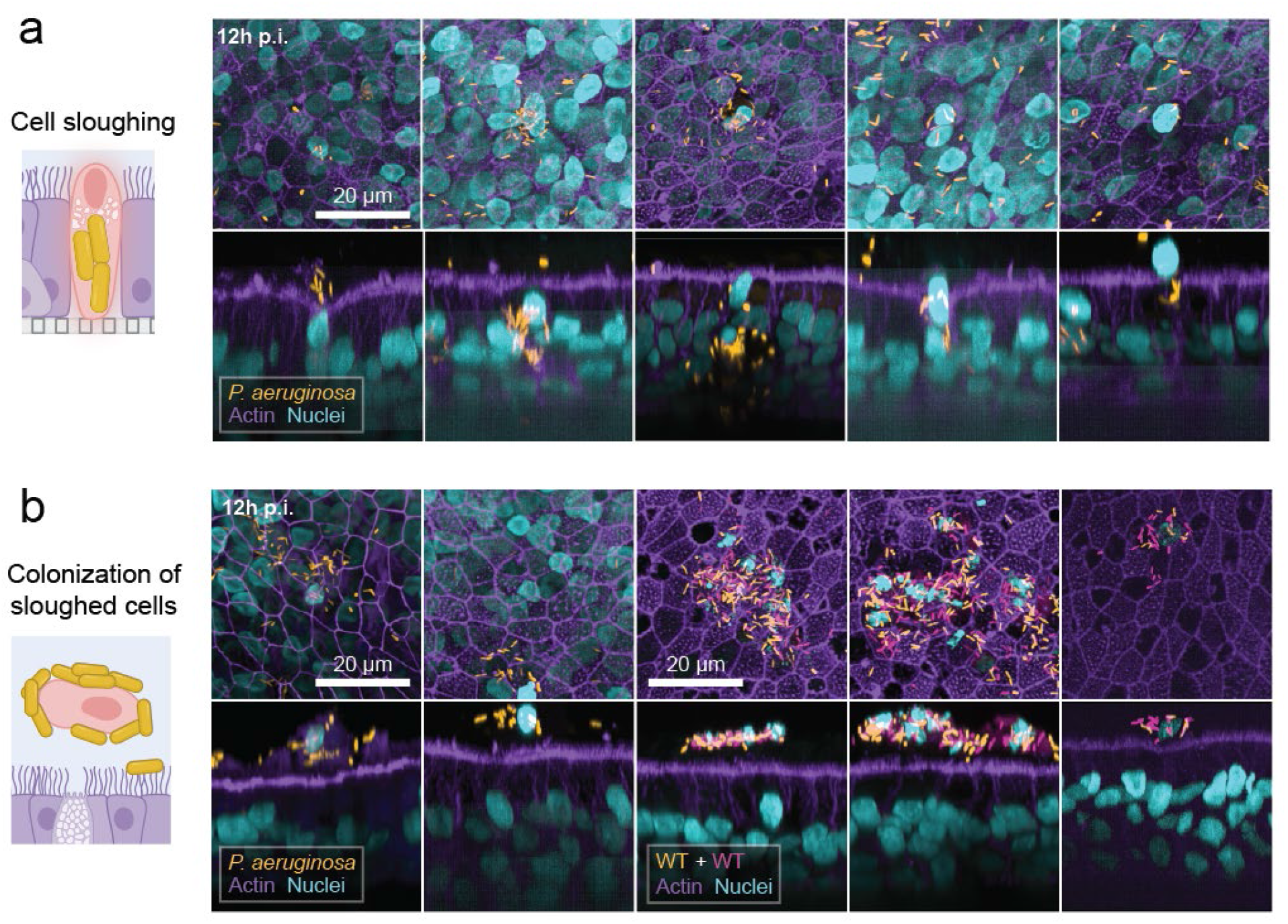
Sloughing of *P. aeruginosa* infected epithelial cells. **a, b**. Representative ICC images (100x) depicting sloughing of epithelial cells infected with *P. aeruginosa* (**a**) and colonization of sloughed cells by extracellular bacteria (**b**). Inoculum=10^3^. *n*=3. Staining: bacteria: constitutive chromosomal expression of mNeonGreen, nuclei: DAPI, actin: phalloidin-647.

**Extended data Fig. 7.**
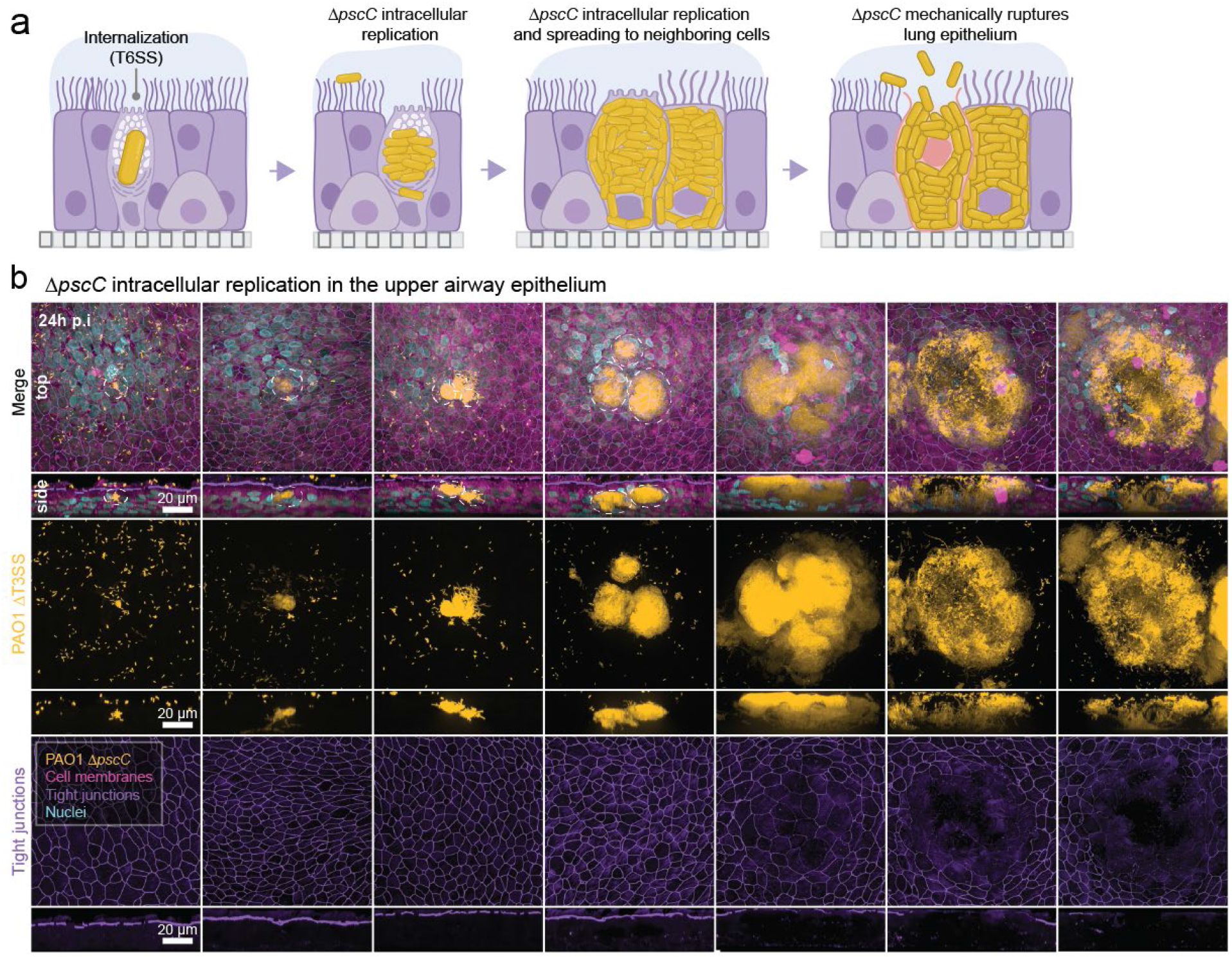
Intracellular replication of *P. aeruginosa* T3SS mutants. **a**, **b**. Schematic (**a**) and 100x ICC images (**b**) of the intracellular replication of a *P. aeruginosa* T3SS mutant (Δ*pscC*) resulting in mechanical cell rupture and release of intracellular bacteria. The progression shown in (**b)** is a compilation of different stages of breaching sites observed in samples 24 h p.i. Inoculum=10^3^. *n*=3. Staining: bacteria: constitutive chromosomal expression of mNeonGreen, nuclei: DAPI, membrane: CellMask, tight junctions: ZO-1 Alexa Fluor 555.

**Extended data Fig. 8.**
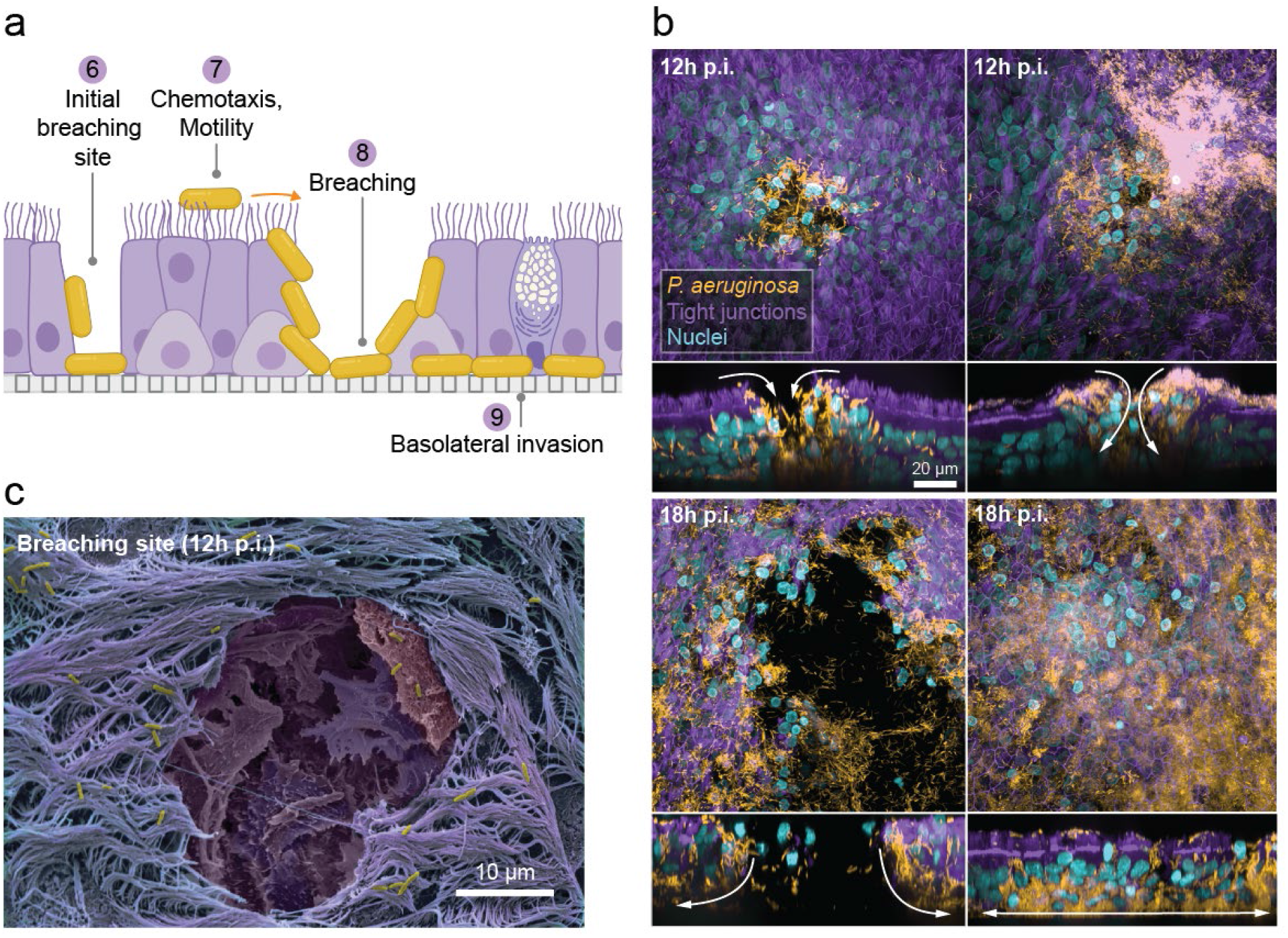
*P. aeruginosa* breaching and basolateral invasion of the upper airway epithelium. **a, b.** Schematic (**a**) and 100x ICC images (**b**) of *P. aeruginosa* invasion of the basolateral side of the lung epithelium after successful establishment of a breaching site. Staining: bacteria: constitutive chromosomal expression of mNeonGreen, nuclei: DAPI, tight junctions: ZO-1 Alexa Fluor 555. **c**. Progression of upper airway infection by *P. aeruginosa* PAO1 wild type recorded by SEM. For all panels: Inoculum=10^3^. *n*=3.

**Extended data Fig. 9.**
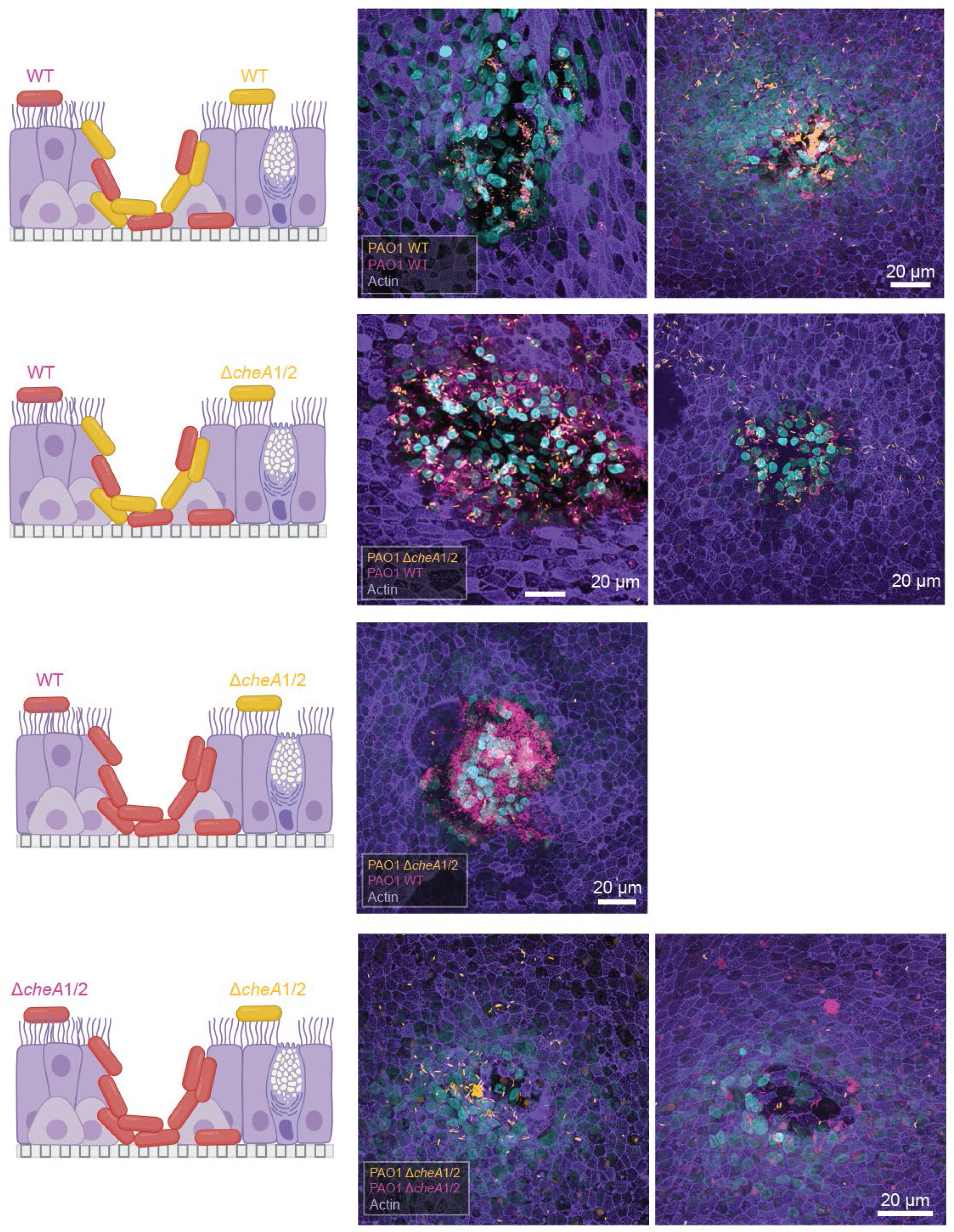
Rapid colonization of breaching sites is driven by chemotaxis. Breaching site colonization in samples infected with a 1:1 mixture of differentially labeled (pink or yellow) wild type and Δ*cheA*1/2 mutant strains. Inoculum=10^3^, *n*=3. Staining: bacteria: constitutive chromosomal expression of mNeonGreen or mCherry, nuclei: DAPI, actin: phalloidin-647.

## Extended data Movies

**Extended data Movie 1. Cilia beating and mucus flow**. Real time recording of beating cilia and mucus flow on the apical surface of the upper airway model. Staining: membrane: CellMask.

**Extended data Movie 2. Tissue regeneration and *P. aeruginosa* infection of injured lung epithelium.** Wound healing after physical tissue damage of uninfected upper airway epithelium (upper panels) and selective *P. aeruginosa* invasion of the insured site upon infection (lower panels). Staining: bacteria: constitutive chromosomal expression of mNeonGreen, membrane: CellMask. Inoculum=10^3^, *n*=3.

**Extended data Movie 3. Increased frequency of apoptotic lung cell events upon infection with *P. aeruginosa*.** Live imaging (25x) of apoptotic events upon infection with *P. aeruginosa*. Staining: apoptotic cells: NucView (caspase-3/7 activity), membrane: CellMask. Inoculum=10^3^, *n*=3.

## Extended data Tables

**Extended data Table 1.** Bacterial strains used in this study. Related to Key Resources Table

| Name | Description | Reference | ID |
| --- | --- | --- | --- |
| ***P. aeruginosa*** |  |  |  |
| PAO1 | Wild-type *Pseudomonas aeruginosa* | <sup>2</sup> | UJP505 |
| $\Delta pilA$ | Unmarked deletion of *pilA* (PA4525) in PAO1 | This study | BJL019 |
| $\Delta fliC$ | Unmarked deletion of *fliC* (PA4332) in PAO1 | <sup>3</sup> | UJP2789 |
| $\Delta pscC$ | Unmarked deletion of *pscC* (PA1716) in PAO1 | <sup>3</sup> | UJP3542 |
| $\Delta pch$ | Unmarked deletion of *pch* (PA5017) in PAO1 | This study | BJL272 |
| $\Delta cyaAB$ | Unmarked deletion of *cyaAB* (PA5272 and PA3217) in PAO1 | This study | UJP4026 |
| $\Delta tssL2$ | Unmarked double deletion of *tssL2* (PA1668) in PAO1 | This study | UJP7885 |
| $\Delta tssM3$ | Unmarked deletion of *tssM3* (PA2361) in PAO1 | This study | UJP7886 |
| $\Delta cheA1-2$ | Unmarked double deletion of *cheA1* and *cheA2* (PA0177 and PA0178) in PAO1 | This study | UJP6929 |
| PA14 | Wild-type *Pseudomonas aeruginosa* | R. Kolter | UJP218 |
| PA Boston 41501 | Wild-type *Pseudomonas aeruginosa* | ATCC 27853 | UJP6789 |
| PAO1 CTX::nGreen | PAO1 with chromosomal integration for constitutive expression of nGreen by the pX2 promoter. | This study | BJL433 |
| $\Delta pilA$ CTX::nGreen | $\Delta pilA$ with chromosomal integration for constitutive expression of nGreen by the pX2 promoter. | This study | UJP7556 |
| $\Delta fliC$ CTX::nGreen | $\Delta fliC$ with chromosomal integration for constitutive expression of nGreen by the pX2 promoter. | This study | UJP7557 |
| $\Delta pscC$ CTX::nGreen | $\Delta pscC$ with chromosomal integration for constitutive expression of nGreen by the pX2 promoter. | This study | UJP6559 |
| $\Delta pch$ CTX::nGreen | $\Delta pch$ with chromosomal integration for constitutive expression of nGreen by the pX2 promoter. | This study | BJL439 |
| $\Delta cyaAB$ CTX::nGreen | $\Delta cyaAB$ with chromosomal integration for constitutive expression of nGreen by the pX2 promoter. | This study | UJP7558 |
| $\Delta tssL2$ CTX::nGreen | $\Delta tssL2$ with chromosomal integration for constitutive expression of nGreen by the pX2 promoter. | This study | UJP7887 |
| $\Delta tssM3$ CTX::nGreen | $\Delta tssM3$ with chromosomal integration for constitutive expression of nGreen by the pX2 promoter. | This study | UJP7888 |
| $\Delta cheA1-2$ CTX::nGreen | $\Delta cheA1-2$ with chromosomal integration for constitutive expression of nGreen by the pX2 promoter. | This study | UJP7789 |
| PA14 CTX::nGreen | PA14 with chromosomal integration for constitutive expression of nGreen by the pX2 promoter. | This study | UJP6561 |
| ***E. coli*** |  |  |  |
| DH5α | *endA1*, *hsdR17*(rK-mK+), *supE44*, *recA1*, *gyrA* (Nalr), *relA1*, $\Delta$(*lacIZYAargF*) U169, *deoR*, Φ80*dlac*Δ(*lacZ*)M15 | - | N/A |
| BL21 lambda DE3 | *E. coli* strain for protein purification: F<sup>-</sup> ompT gal dcm lon hsdSB(rB-mB-) $\lambda$(DE3 [*lacI* *lacUV5*-T7 gene 1 ind1 sam7 nin5]) | New England Biolabs | Cat#C2527 |

**Extended data Table 2.** Key resources used in this study.

| REAGENT or RESOURCE | SOURCE | IDENTIFIER |
| --- | --- | --- |
| **Antibodies** |  |  |
| Mouse anti mucin 5AC 1:200 | abcam | Cat # Ab3649 |
| Rabbit anti acetyl-alpha-Tubulin (Lys40) Alexa 647 conjugate 1:50 | Cell Signaling | Cat # 81502 |
| 555 ZO-1 mouse Alexa 555 conjugate 1:200 | Invitrogen | Cat # MA3-39100-A555 |
| Chicken anti Ki67 1:1000 | abcam | Cat # Ab254123 |
| Mouse anti p63 1:50 | abcam | Cat # Ab735 |
| Rat anti SCGB1A1(uteroglobin) 1:50 | R&D System | Cat # MAB4218 |
| Mouse anti MUC5AC (Clone 917) 1:400 | Novus Biologicals | Cat# NBP2-54448AF488 |
| Rabbit anti P63-PE, (Clone D2K8X) 1:100 | Cell Signaling | Cat# 56687S |
| Rabbit anti alpha tubulin AF647, (Clone EPR16772) 1:50 | abcam | Cat# ab218591 |
| Cy3 donkey anti chicken 1:2000 | Jackson Immuno Research | Cat # 703-166-155 |
| Cy3 donkey anti rat 1:2000 | Jackson Immuno Research | Cat # 712-166-150 |
| 488 donkey anti mouse 1:800 | Invitrogen | Cat # A32766 |
| Cy3 donkey anti-mouse 1:2000 | Jackson Immuno Research | Cat # 715-165-151 |
| **Bacterial and Virus Strains** |  |  |
| *A detailed strain list can be found in Supplemental Table S1* |  |  |
| **Chemicals, Peptides, and Recombinant Proteins** |  |  |
| PneumaCult™-Ex Plus Medium | Stem Cell Technologies | Cat # 05040 |
| PneumaCult-ALI KIT | Stem Cell Technologies | Cat # 05001 |
| Hydrocortisone (medium supplement) | StemCell Technologies | Cat # 07926 |
| Heparin Solution (medium supplement) | StemCell Technologies | Cat # 07980 |
| Costar 6.5 mm Transwell®, 0.4 µm Pore Polyester Membrane Inserts | StemCell Technologies | Cat # 38024 |
| 75 cm<sup>2</sup> Cell culture flasks | Falcon | Cat # 353136 |
| HBSS | Gibco | Cat # 14025050 |
| Formalin | Sigma-Aldrich | Cat # HT5011-1CS |
| Mowiol | Sigma-Aldrich | Cat # 81381 |
| Glass slides Super Frost | Fisher scientific | Cat # 16291541 |
| cover slips #1.5 | VWR | Cat # 631-0147 |
| Triton 100 | Sigma-Aldrich | Cat # T8787 |
| Sodium Azide | Sigma-Aldrich | Cat # S2002 |
| FCS | ThermoFisher | Cat # 10082147 |
| Trypsin | Gibco | Cat # 15400-054 |
| Alexa 647 Phalloidin 1:250 | Molecular Probes | Cat # A22287 |
| Alexa 546 Phalloidin 1:500 | Molecular Probes | Cat # A22283 |
| 647CellMask DeepRed fix:1:10 000 | ThermoFisher | Cat # A57245 |
| DAPI 1:1000 | Sigma-Aldrich | Cat # D9542 |
| NucView<sup>R</sup> 488 | Biotium | Cat # 10403-T |
| AccuMax | Sigma-Aldrich | Cat # A7089 |
| Animal Component-Free Cell Dissociation Kit | StemCell Technologies | Cat # 05426 |
| MycoStrip | InvivoGen | Cat # rep-mys-10 |
| **Experimental Models: Cell Lines** |  |  |
| NHBE stem cells | Lonza | CC-2540S, donor 36311, Lot 18TL215671 |

## Extended Methods

### Contact for reagent and resource sharing

Further information and requests for resources should be directed to the Lead Contact: Urs Jenal.

### Strains, plasmids and growth media

Strains used in this study are listed in Extended data Table 1. *P. aeruginosa* PAO1 and mutants were grown at 37°C under shaking conditions.

### Molecular biology procedures

Fluorescent *P. aeruginosa* strains were generated using an integrative plasmid encoding mNeonGreen under control of a constitutive pX2 promoter. The strains were transformed by electroporation and selected on LB Tet 100 μg/mL agar plates.

Deletions of *tssL2* (T6SS-H2) and *tssM3* (T6SS-H3) in *P. aeruginosa* strain PAO1 were generated by two-step allelic exchange^4^. We used the integrative plasmid pEX18 (TetR) in which we cloned the regions flanking the target gene to delete (c.a. 700bp). The 2 flanking regions were amplified individually as described above, then fused in a second step by overlap extension PCR (SOE-PCR). *P. aeruginosa* was then transformed by mating with a donor *E. coli* SM10. *P. aeruginosa* mutants were selected by sucrose counter-selection on 5% sucrose LB agar plates. Tetracycline sensitive clones were selected by pick and patching clones on LB and LB Tet 100 mg/mL agar plates. Mutants were validated by colony PCR.

### Cell culture

Normal Human Bronchial Epithelial (NHBE) cells (Lonza) were expanded and differentiated as described by StemCell Technologies (stemcell.com). In short, frozen passage-1 NHBE stocks were thawed and cultured in T75-flask using PneumaCult-Ex Plus medium (Stemcell Technologies) at 37°C in an atmosphere of 5% CO_2_ and 95% relative humidity. Once at 60– 80% confluent, cells were dissociated using ACF Enzymatic Dissociation Solution (Stemcell Technologies) and seeded on 6.5 mm Transwell inserts (Corning) with a 0.4 µm pore size. Cells were seeded at 3,3 x 10^4^ cells/insert and cultured for 2-3 days at liquid-liquid interface in PneumaCult-Ex Plus medium till full confluency was reached. Subsequently, the apical media was removed and the basal media replaced with PneumaCult-ALI medium (Stemcell Technologies). The medium was changed every second or third day and the apical surface washed with HBSS (without Ca^2+^, Mg^2+^) once per week from age 25 onwards. The cells were cultured for a minimum of 25 days post airlift before experiments were performed.

### Lung tissue infection and CFU assay

*P. aeruginosa* was grown to stationary phase in synthetic cystic fibrosis medium (SCFM)^1^ liquid cultures at 37°C under shaking conditions. The cultures were back-diluted 1:100 in SCFM and growth for 3-4 hours at 37°C under shaking conditions. Bacteria were washed with and adjusted in PBS to OD600 = 1.0-0.001. Lung tissues were infected with 2µl of the adjusted suspension with to the inoculum size indicated, 10^1-10^6 respectively. For the meropenem treatment experiment, the antibiotic was added to the basolateral compartment at 12 hours post infection and subsequently removed at 24 hours post infection.

To assess bacterial replication on the lung tissue surface, the lung tissue and membrane were cut from the insert and dissociated with 2 mm glass beads in ice-cold PBS by 30 sec vortexing. The PBS suspension was serial-diluted and plates on LB agar. Agar plates were grown overnight at 37°C and CFU were counted. In parallel, tissue breaching was assessed by sampling the basolateral compartment, and plating the serial dilutions. CFUs were counted manually and reported in CFU/ml.

### Viability assay

Tissue viability was assessed according to the manual of the Cytotoxicity Detection Kit^PLUS^ (CYTODET-RO4744934001, Sigma). In short, 100 µl medium was collected from the basolateral compartment and mixed with 100 µl of reaction buffer, containing dye solution and lyophilized catalyst (45:1). The samples were incubated for max. 30 min at RT in the dark. The reaction was stopped by addition of 50 µl stop solution and the absorbance at 500 nm was measured using an Epoch Microplate Spectrophotometer (Biotek). For each experiment, an internal positive control was included where a lung tissue was treated for 24 hours with lysis solution (included in kit) and the measured absorbance value was set to 100%.

### Flow cytometry

Cell viability upon infection and cell type populations were assessed with flow cytometry. To this end, the tissues were dissociated with AccuMax (Sigma, A7089) on the apical and basolateral for 30 min at RT. Resuspended single cells were collected in FACS buffer (PBS 2% FCS 2mM EDTA) and stained with Zombie NIR™ Fixable Viability kit (Biolegend, 423105) 1:1000 in PBS for 15 minutes at RT and subsequently washed with PBS. The cells were treated with Foxp3 Fixation/Permeabilization working solution (00-5523-00, ThermoFisher) for 30min at RT and washed twice with 1X permeabilization buffer (1:10 in ddH_2_0). For cell type population analysis, the cells were treated for 15 min at RT with Human TruStain FcX™ (Fc Receptor Blocking Solution, 422301, Biolegend) 1:10 in Permeabilization buffer. Samples were directly mixed with 2x antibody master mix containing 1:400 anti-MUC5AC (Clone 917, Catalog# NBP2-54448AF488), 1:100 anti-P63-PE, (Clone D2K8X, Catalog# 56687S) and 1:50 anti-alpha tubulin AF647, (Clone EPR16772, Catalog# ab218591) and incubate for 30 minutes at 4°C. The cells were washed twice and stored in FACS buffer till analysis. The samples were analyzed by BD LSR Fortessa (Cell type populations) or BD Canto II (cell viability). Data was processed with BD FACSDiva software.

### Cilia beating frequency measurements

Live cilia beating movies were recorded using a Nikon CSU-W1 microscope, equipped with a 2x Hamamatsu ORCA-Fusion camera (sCMOS) and Plan Fluor 4x/0.13 and Plan Apo Lambda 10x/0.45 objectives, at 37°C in an atmosphere of 5% CO_2_ and 95% relative humidity. The microscope was set-up in wide field mode, with a 4x air objective and a total of 512 frames were recorded at a frame rate of 200 fps. The videos were processed with the Cilia-X software (Epithelix) to measure cilia beating frequency and the percentage of regions showing actively beating cilia (threshold: >3.5 Hz).

### TEER measurements

Transepithelial electrical resistance (TEER) values of the lung tissues at distinct time point post airlift or infection were measured with EVOM3 Epithelial Volt Ohm Meter using the STX2-Plus Electrode (World Precision Instruments). The tissues were mucus washed and submerged in pre-warmed HBSS (350 µL apical and 1 ml basolateral) for the duration of the measurement. TEER values of the empty inserts were subtracted from the measured values and the reported value (TEER_REPORTED_, in units of Ω.cm^2^) was calculated as:

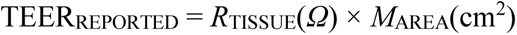

Post measurement, HBSS was removed and tissues were sacrificed or further cultured as described above.

### Time-lapse live imaging

Lung tissues were stained with 2 µg/mL CellMask™ Deep Red Plasma Membrane Stain (ThermoFisher) in the basal compartment for 24-48 hours prior infection and imaging, and imaged in 24-well plates. To visualize apoptotic cells, after CellMask™ Deep Red staining, the lung tissue was stained with 5 µM NucView® 488 Caspase-3 Substrate (Biotium) for 40 min, then imaged in a Delta T Transwell Insert Adapter (Bioptechs).

Samples were imaged using a Nikon CSU-W1 equipped with a 2x Hamamatsu ORCA-Fusion camera (sCMOS) and CFP Plan Apo Lambda 10x/0.45 or CFI Plan Apo Lambda S 25x/1.05 silicon objective. Images were stored and analyzed using OMERO.

### Immunocytochemistry

At the indicated tissue age or experimental time point, samples were washed at the basolateral side with HBSS and subsequently fixed by addition of 10% formalin to the apical and basal compartment, for 45 min at RT. Samples were washed once with HBSS containing 0.05% NaN_3_ and stored at 4°C in this buffer until further processing. The membranes were cut from the inserts and processed for ICC or histology. For ICC, the tissues were permeabilized for 20 min at RT with HBBS 0.5 % Triton. Then, washed once with HBSS with NaN_3_ 0,05% and stained with the respective primary and secondary antibodies and/or dyes (see key resource table). The samples were embedded in Mowiol and cured for >48 hours at RT before imaging. ICC samples were imaged with Olympus SpinD (CSU-W1) equipped with a Hamamatsu ORCA-Fusion (sCMOS) camera and UPL X APO 10x/0.4 and 30x/1.05 objectives or Nikon CSU-W1 equipped with an 2x Hamamatsu ORCA-Fusion camera (sCMOS) and CFP Plan Apo Lambda 100x/1.45 objectives. Image analysis and quantification was performed with Imaris Imaging Analysis Software or NIS-Elements General Analysis software.

### Histology

Inserts were fixed as described for immunocytochemistry samples. Insert’s membranes were embedded in paraffin, sectioned in 4 µm thick slices and stained with hematoxylin and eosin using an automated stainer. Histology samples were imaged with Zeiss AxioScan.Z1 Slidescanner equipped with an 3CCD Hitachi HV-F202SCL camera and Plan Apochromat 40x/0.95 objective.

### Scanning electron microscopy

At time-point indicated, the apical and basolateral compartments of the sample was washed with HBSS. HBSS was removed and replaced with the fixative, 4% formaldehyde + 1% glutaraldehyde in PBS. The sample was incubated for 24 hours at RT and subsequently washed twice with ddH_2_O. The sample was dehydrated with different 10 min incubation steps in 30%, 50%, 70% and 90% EtOH (EMS, 15055) solutions in ddH_2_O and then incubated twice of 15 min in 100% EtOH. The samples were incubated for 20 min in a 1:2, followed by a 2:1 solution of Hexamethyldisilazane (HMDS, Sigma, 440191) and 100% EtOH. Next, the sample was incubated twice for 20 min in pure Hexamethyldisilazane and subsequently incubated overnight in Hexamethyldisilazane while letting the solution evaporate. Once dry, the membrane was cut from the insert and fixed onto a sample holder with a conductive carbon sticker (Ted Pella, 77852). The sample was metal coated with Quorum SC7620 with platinum/Au alloy for 30 seconds.

SEM imaging was performed with a FEI Versa 3D microscope (Thermo Fisher Scientific) or Philips XL30 ESEM microscope (modified by RemX GmbH, Germany) under high vacuum at an acceleration voltage of 5KV. The images were collected using the SE Everhard Thornley Detector at different magnifications.

### Statistics and reproducibility

Statistical data analysis was performed in GraphPad Prism 9. Unless otherwise indicated, all experiments were performed in three independent biological replicates. Measurements were taken from distinct samples, only for the CBF the same sample was repeatedly measured over time. All microscopy images shown are representative of experiments in which a minimum of three Transwell lung tissues were surveyed with similar results. For quantitative microscopy data, three independent experiments analyzing at least 15 images per sample; mean values from independent experiments were used. In all graphs, data are presented as mean ± standard deviation (SD). For box plots: center line, median; box limits, upper and lower quartiles; whiskers, min-max. Information about specific statistical tests in each analysis can be found in the figure legends. For unpaired t tests, normal distribution was not tested but assumed. Exact *P* values of statistical testing can be found in source data files. No statistical methods were used to pre-determine sample sizes, but our sample sizes are similar to those reported in previous publications.

## References

1. Giantsou, E. & Manolas, K. I. Superinfections in *Pseudomonas aeruginosa* ventilator-associated pneumonia. Minerva Anestesiol 77, 964–70 (2011).

2. Happel, K. I., Nelson, S. & Summer, W. The Lung in Sepsis: Fueling the Fire. Am J Medical Sci 328, 230–237 (2004).

3. Meyer, N. J., Gattinoni, L. & Calfee, C. S. Acute respiratory distress syndrome. Lancet 398, 622–637 (2021).

4. Berube, B. J., Rangel, S. M. & Hauser, A. R. *Pseudomonas aeruginosa*: breaking down barriers. Curr Genet 62, 109–113 (2016).

5. Gellatly, S. L. & Hancock, R. E. W. *Pseudomonas aeruginosa*: new insights into pathogenesis and host defenses. Pathog Dis 67, 159–173 (2013).

6. Jurado-Martín, I., Sainz-Mejías, M. & McClean, S. *Pseudomonas aeruginosa*: An Audacious Pathogen with an Adaptable Arsenal of Virulence Factors. Int J Mol Sci 22, 3128 (2021).

7. Parker, D. & Prince, A. Innate Immunity in the Respiratory Epithelium. Am J Resp Cell Mol 45, 189–201 (2011).

8. Tsang, K. et al. Interaction of *Pseudomonas aeruginosa* with human respiratory mucosa in vitro. Eur Respir J 7, 1746–1753 (1994).

9. Schwarzer, C., Fischer, H. & Machen, T. E. Chemotaxis and Binding of *Pseudomonas aeruginosa* to Scratch-Wounded Human Cystic Fibrosis Airway Epithelial Cells. Plos One 11, e0150109 (2016).

10. Tramper-Stranders, G. A., et al. Initial *Pseudomonas aeruginosa* infection in patients with cystic fibrosis: characteristics of eradicated and persistent isolates. Clin Microbiol Infec 18, 567–574 (2012).

11. Bustamante-Marin, X. M. & Ostrowski, L. E. Cilia and Mucociliary Clearance. Csh Perspect Biol 9, a028241 (2017).

12. Carvajal, A. & Pérez, P. Epidemiology of Respiratory Infections. *In:* Pediatric Respiratory Diseases: A Comprehensive Textbook. (Springer International Publishing, 2020).

13. Hiemstra, P. S., McCray, P. B. & Bals, R. The innate immune function of airway epithelial cells in inflammatory lung disease. Eur Respir J 45, 1150–1162 (2015).

14. Organization, W. H. World Health Organization Global Tuberculosis Program. Global Tuberculosis Report 2020. https://www.who.int/publications/i/item/9789240037021 (2021).

15. Almagro, P., et al. *Pseudomonas aeruginosa* and Mortality after Hospital Admission for Chronic Obstructive Pulmonary Disease. Respiration 84, 36–43 (2012).

16. Engel, J. & Balachandran, P. Role of *Pseudomonas aeruginosa* type III effectors in disease. Curr Opin Microbiol 12, 61–66 (2009).

17. Fleiszig, S. M. et al. Epithelial cell polarity affects susceptibility to Pseudomonas aeruginosa invasion and cytotoxicity. Infect Immun 65, 2861–2867 (1997).

18. Engel, J. & Eran, Y. Subversion of Mucosal Barrier Polarity by *Pseudomonas aeruginosa*. Front Microbiol 2, 114 (2011).

19. Golovkine, G., et al. *Pseudomonas aeruginosa* Transmigrates at Epithelial Cell-Cell Junctions, Exploiting Sites of Cell Division and Senescent Cell Extrusion. PLoS Pathog 12, e1005377 (2016).

20. Kierbel, A., et al. *Pseudomonas aeruginosa* exploits a PIP3-dependent pathway to transform apical into basolateral membrane. J Cell Biology 177, 21–27 (2007).

21. Kierbel, A., Gassama-Diagne, A., Mostov, K. & Engel, J. N. The Phosphoinositol-3-Kinase–Protein Kinase B/Akt Pathway Is Critical for *Pseudomonas aeruginosa* Strain PAK Internalization. Mol Biol Cell 16, 2577–2585 (2005).

22. Lopes, S. F. et al. Primary and Immortalized Human Respiratory Cells Display Different Patterns of Cytotoxicity and Cytokine Release upon Exposure to Deoxynivalenol, Nivalenol and Fusarenon-X. Toxins 9, 337 (2017).

23. Barron, S. L., Saez, J. & Owens, R. M. In Vitro Models for Studying Respiratory Host– Pathogen Interactions. Adv Biology 5, 2000624 (2021).

24. Seok, J. et al. Genomic responses in mouse models poorly mimic human inflammatory diseases. Proc National Acad Sci 110, 3507–3512 (2013).

25. Worp, H. B. van der et al. Can Animal Models of Disease Reliably Inform Human Studies? PLoS Med 7, e1000245 (2010).

26. Uhl, E. W. & Warner, N. J. Mouse Models as Predictors of Human Responses: Evolutionary Medicine. Curr Pathobiology Reports 3, 219–223 (2015).

27. Chia, S. P. S., Kong, S. L. Y., Pang, J. K. S. & Soh, B.-S. 3D Human Organoids: The Next “Viral” Model for the Molecular Basis of Infectious Diseases. Biomed 10, 1541 (2022).

28. Sato, T. et al. Long-term Expansion of Epithelial Organoids From Human Colon, Adenoma, Adenocarcinoma, and Barrett’s Epithelium. Gastroenterology 141, 1762–1772 (2011).

29. Lee, S. H. et al. Tumor Evolution and Drug Response in Patient-Derived Organoid Models of Bladder Cancer. Cell 173, 515–528.e17 (2018).

30. Loomans, C. J. M. et al. Expansion of Adult Human Pancreatic Tissue Yields Organoids Harboring Progenitor Cells with Endocrine Differentiation Potential. Stem Cell Rep 10, 712– 724 (2018).

31. Guan, Y. et al. A human multi-lineage hepatic organoid model for liver fibrosis. Nat Commun 12, 6138 (2021).

32. Rock, J. R. et al. Basal cells as stem cells of the mouse trachea and human airway epithelium. Proc National Acad Sci 106, 12771–12775 (2009).

33. Youk, J. et al. Three-Dimensional Human Alveolar Stem Cell Culture Models Reveal Infection Response to SARS-CoV-2. Cell Stem Cell 27, 905–919.e10 (2020).

34. Schwank, G. et al. Functional Repair of CFTR by CRISPR/Cas9 in Intestinal Stem Cell Organoids of Cystic Fibrosis Patients. Cell Stem Cell 13, 653–658 (2013).

35. Sachs, N. et al. Long-term expanding human airway organoids for disease modeling. EMBO J 38, (2019).

36. Han, Y. et al. Identification of SARS-CoV-2 inhibitors using lung and colonic organoids. Nature 589, 270–275 (2021).

37. Heo, I. et al. Modelling *Cryptosporidium* infection in human small intestinal and lung organoids. Nat Microbiol 3, 814–823 (2018).

38. García, S. R. et al. Novel dynamics of human mucociliary differentiation revealed by single-cell RNA sequencing of nasal epithelial cultures. Development 146, dev.177428 (2019).

39. Fahy, J. V. & Dickey, B. F. Airway Mucus Function and Dysfunction. New Engl J Medicine 363, 2233–2247 (2010).

40. Liao, C., Huang, X., Wang, Q., Yao, D. & Lu, W. Virulence Factors of *Pseudomonas aeruginosa* and Antivirulence Strategies to Combat Its Drug Resistance. Front Cell Infect Mi 12, 926758 (2022).

41. Williams, P. & Cámara, M. Quorum sensing and environmental adaptation in *Pseudomonas aeruginosa*: a tale of regulatory networks and multifunctional signal molecules. Curr Opin Microbiol 12, 182–191 (2009).

42. Jimenez, P. N. et al. The Multiple Signaling Systems Regulating Virulence in *Pseudomonas aeruginosa*. Microbiology and Molecular Biology Reviews 76, 46–65 (2012).

43. Persat, A., Inclán, Y. F., Engel, J. N., Stone, H. A. & Gitai, Z. Type IV pili mechanochemically regulate virulence factors in *Pseudomonas aeruginosa*. Proceedings of the National Academy of Sciences 112, 7563–7568 (2015).

44. Klockgether, J. & Tümmler, B. Recent advances in understanding *Pseudomonas aeruginosa* as a pathogen. F1000Research 6, 1261 (2017).

45. Bleves, S. et al. Protein secretion systems in Pseudomonas aeruginosa: A wealth of pathogenic weapons. Int J Med Microbiol 300, 534–543 (2010).

46. O’Toole, G. A. & Kolter, R. Flagellar and twitching motility are necessary for *Pseudomonas aeruginosa* biofilm development. Molecular Microbiology 30, 295–304 (1998).

47. Giltner, C. L., Nguyen, Y. & Burrows, L. L. Type IV Pilin Proteins: Versatile Molecular Modules. Microbiology and Molecular Biology Reviews 76, 740–772 (2012).

48. Laventie, B.-J. et al. A Surface-Induced Asymmetric Program Promotes Tissue Colonization by *Pseudomonas aeruginosa*. Cell Host & Microbe 25, 140–152.e6 (2019).

49. Luo, Y. et al. A Hierarchical Cascade of Second Messengers Regulates Pseudomonas aeruginosa Surface Behaviors. mBio 6, e02456–14 (2015).

50. Kazmierczak, B. I., Schniederberend, M. & Jain, R. Cross-regulation of *Pseudomonas* motility systems: the intimate relationship between flagella, pili and virulence. Current Opinion in Microbiology 28, 78–82 (2015).

51. Valentini, M. & Filloux, A. Multiple Roles of c-di-GMP Signaling in Bacterial Pathogenesis. Annual Review of Microbiology 73, 387–406 (2019).

52. Laventie, B.-J. & Jenal, U. Surface Sensing and Adaptation in Bacteria. Annu Rev Microbiol 74, 735–760 (2020).

53. Palmer, K. L., Aye, L. M. & Whiteley, M. Nutritional Cues Control *Pseudomonas aeruginosa* Multicellular Behavior in Cystic Fibrosis Sputum. J Bacteriol 189, 8079–8087 (2007).

54. Kuek, L. E. & Lee, R. J. First contact: the role of respiratory cilia in host-pathogen interactions in the airways. Am J Physiol-lung C 319, L603–L619 (2020).

55. Widdicombe, J. H. & Wine, J. J. Airway Gland Structure and Function. Physiol Rev 95, 1241–1319 (2015).

56. Sana, T. G. et al. Internalization of *Pseudomonas aeruginosa* Strain PAO1 into Epithelial Cells Is Promoted by Interaction of a T6SS Effector with the Microtubule Network. mBio 6, e00712–15 (2015).

57. Sana, T. G. et al. The Second Type VI Secretion System of *Pseudomonas aeruginosa* Strain PAO1 Is Regulated by Quorum Sensing and Fur and Modulates Internalization in Epithelial Cells. J Biol Chem 287, 27095–27105 (2012).

58. Hogan, B. L. M. et al. Repair and Regeneration of the Respiratory System: Complexity, Plasticity, and Mechanisms of Lung Stem Cell Function. Cell Stem Cell 15, 123–138 (2014).

59. Rossy, T. et al. *Pseudomonas aeruginosa* contracts mucus to rapidly form biofilms in tissue-engineered human airways. Biorxiv 2022.05.26.493615 (2022) doi:10.1101/2022.05.26.493615.

60. McDole, J. R. et al. Goblet cells deliver luminal antigen to CD103+ dendritic cells in the small intestine. Nature 483, 345–349 (2012).

61. Jiang, F., Waterfield, N. R., Yang, J., Yang, G. & Jin, Q. A *Pseudomonas aeruginosa* Type VI Secretion Phospholipase D Effector Targets Both Prokaryotic and Eukaryotic Cells. Cell Host Microbe 15, 600–610 (2014).

62. Rajan, S., et al. *Pseudomonas aeruginosa* Induction of Apoptosis in Respiratory Epithelial Cells. Am J Resp Cell Mol 23, 304–312 (2000).

63. Yamaguchi, T. & Yamada, H. Role of Mechanical Injury on Airway Surface in the Pathogenesis of *Pseudomonas aeruginosa*. Am Rev Respir Dis 144, 1147–1152 (1991).

64. Heiniger, R. W., Winther-Larsen, H. C., Pickles, R. J., Koomey, M. & Wolfgang, M. C. Infection of human mucosal tissue by *Pseudomonas aeruginosa* requires sequential and mutually dependent virulence factors and a novel pilus-associated adhesin. Cellular microbiology 12, 1158–1173 (2010).

65. Kumar, N. G. et al. Pseudomonas aeruginosa Can Diversify after Host Cell Invasion to Establish Multiple Intracellular Niches. mBio 13, e02742–22 (2022).

66. Chastre, J. et al. Safety, efficacy, and pharmacokinetics of gremubamab (MEDI3902), an anti-*Pseudomonas aeruginosa* bispecific human monoclonal antibody, in *P. aeruginosa*-colonised, mechanically ventilated intensive care unit patients: a randomised controlled trial. Crit Care 26, 355 (2022).

67. Hotinger, J. A. & May, A. E. Antibodies Inhibiting the Type III Secretion System of Gram-Negative Pathogenic Bacteria. Antibodies 9, 35 (2020).

68. Jain, M. et al. Type III Secretion Phenotypes of *Pseudomonas aeruginosa* Strains Change during Infection of Individuals with Cystic Fibrosis. J Clin Microbiol 42, 5229–5237 (2004).

69. Huus, K. E. et al. Clinical Isolates of *Pseudomonas aeruginosa* from Chronically Infected Cystic Fibrosis Patients Fail To Activate the Inflammasome during Both Stable Infection and Pulmonary Exacerbation. J Immunol 196, 3097–3108 (2016).

70. Rossi, E. et al. Pseudomonas aeruginosa adaptation and evolution in patients with cystic fibrosis. Nat Rev Microbiol 1–12 (2020) doi:10.1038/s41579-020-00477-5.

71. Osan, J. et al. Goblet Cell Hyperplasia Increases SARS-CoV-2 Infection in Chronic Obstructive Pulmonary Disease. Microbiol Spectr 10, e00459–22 (2022).

72. Adam, D. et al. Cystic fibrosis airway epithelium remodelling: involvement of inflammation. J. Pathol. 235, 408–419 (2015).

73. Jeffery, P. K. Comparison of the Structural and Inflammatory Features of COPD and Asthma Giles F. Filley Lecture. Chest 117, 251S–260S (2000).

74. Dovey, M. et al. Ultrastructural morphology of the lung in cystic fibrosis. J Submicr Cytol Path 21, 521–34 (1989).

75. Jeffery, P. K. Remodeling and Inflammation of Bronchi in Asthma and Chronic Obstructive Pulmonary Disease. Proc Am Thorac Soc 1, 176–183 (2004).

76. Ackermann, M. et al. Self-destructive cooperation mediated by phenotypic noise. Nature 454, 987–990 (2008).

77. Diard, M. et al. Stabilization of cooperative virulence by the expression of an avirulent phenotype. Nature 494, 353–356 (2013).

78. Kotte, O., Volkmer, B., Radzikowski, J. L. & Heinemann, M. Phenotypic bistability in *Escherichia coli*’s central carbon metabolism. Molecular Systems Biology 10, 736 (2014).

79. Basan, M. et al. A universal trade-off between growth and lag in fluctuating environments. Nature 584, 470–474 (2020).

80. Bakshi, S. et al. Tracking bacterial lineages in complex and dynamic environments with applications for growth control and persistence. Nat Microbiol 6, 783–791 (2021).

81. Balaban, N. Q., Merrin, J., Chait, R., Kowalik, L. & Leibler, S. Bacterial persistence as a phenotypic switch. Science 305, 1622–1625 (2004).

82. Arnoldini, M. et al. Bistable Expression of Virulence Genes in Salmonella Leads to the Formation of an Antibiotic-Tolerant Subpopulation. PLoS Biol 12, e1001928 (2014).

83. Manina, G., Griego, A., Singh, L. K., McKinney, J. D. & Dhar, N. Preexisting variation in DNA damage response predicts the fate of single mycobacteria under stress. EMBO J 38, e101876 (2019).

84. Keilberg, D., Zavros, Y., Shepherd, B., Salama, N. R. & Ottemann, K. M. Spatial and Temporal Shifts in Bacterial Biogeography and Gland Occupation during the Development of a Chronic Infection. mBio 7, e01705–16 (2016).

85. Fung, C. et al. High-resolution mapping reveals that microniches in the gastric glands control *Helicobacter pylori* colonization of the stomach. PLoS Biol 17, e3000231 (2019).

86. Garvis, S., et al. *Caenorhabditis elegans* Semi-Automated Liquid Screen Reveals a Specialized Role for the Chemotaxis Gene cheB2 in *Pseudomonas aeruginosa* Virulence. PLoS Pathogens 5, e1000540 (2009).

87. Laganenka, L. et al. Chemotaxis and autoinducer-2 signalling mediate colonization and contribute to co-existence of *Escherichia coli* strains in the murine gut. Nat Microbiol 8, 204– 217 (2023).

88. Cooper, K. G. et al. Regulatory protein HilD stimulates *Salmonella typhimurium* invasiveness by promoting smooth swimming via the methyl-accepting chemotaxis protein McpC. Nat Commun 12, 348 (2021).

89. Lane, M. C. et al. Role of Motility in the Colonization of Uropathogenic *Escherichia coli* in the Urinary Tract. Infect Immun 73, 7644–7656 (2005).

## References

1. Huus, K. E. et al. Clinical Isolates of *Pseudomonas aeruginosa* from Chronically Infected Cystic Fibrosis Patients Fail To Activate the Inflammasome during Both Stable Infection and Pulmonary Exacerbation. J Immunol 196, 3097–3108 (2016).

2. Malone, J. G. et al. YfiBNR Mediates Cyclic di-GMP Dependent Small Colony Variant Formation and Persistence in *Pseudomonas aeruginosa*. PLoS Pathogens 6, e1000804 (2010).

3. Laventie, B.-J. et al. A Surface-Induced Asymmetric Program Promotes Tissue Colonization by *Pseudomonas aeruginosa*. Cell Host & Microbe 25, 140–152.e6 (2019).

4. Broder, U. N., Jaeger, T. & Jenal, U. LadS is a calcium-responsive kinase that induces acute-to-chronic virulence switch in *Pseudomonas aeruginosa*. Nature Microbiology 2, 1– 11 (2016).

